# scUnique: Measuring Ongoing Chromosomal Instability

**DOI:** 10.64898/2026.04.28.721240

**Authors:** Michael P. Schneider, Shadi Shafighi, Amy E. Cullen, Akane Ota, Thomas Bradley, Dominique-Laurent Couturier, Philip S. Smith, Anna M. Piskorz, James D. Brenton, Florian Markowetz

## Abstract

Chromosomal instability is a hallmark of cancer that can drive tumour heterogeneity and treatment resistance. However, scalable methods to measure the dynamics of chromosomal instability are lacking and thus historic genomic scars can be hard to distinguish from markers of currently active mutational processes. Here, we introduce *scUnique*, a novel approach to measure ongoing chromosomal instability from single-cell whole genome sequencing (scWGS) data. *scUnique* performs phylogeny-aware joint segmentation to refine single-cell copy number profiles and identifies recent copy number aberrations as statistically supported changes on the leaves of the inferred phylogenetic tree. This yields a per-cell measure of ongoing chromosomal instability at genome-wide resolution. We validate *scUnique* in (i) a comprehensive benchmarking study, (ii) in CRISPR-Cas-based experimental systems, and (iii) single-cell derived clones of a well-characterised ovarian model. We show that *scUnique* distinguishes ongoing HRD and FBI mutational processes. These results show that *scUnique* provides quantitative, scalable, genome-level information about ongoing chromosomal instability, not available in previous studies relying on bulk DNA-sequencing or single-cell imaging. In the future, these improved measurements could refine the understanding of mechanisms of chromosomal instability and lead to dynamic biomarkers for improved treatment decisions.

## 1 Introduction

Chromosomal instability (CIN) is a dynamic process that causes the repeated generation of genomic alterations from the kilobase scale to whole chromosomes. Different mutational processes contribute to these alterations, including homologous recombination deficiency (HRD) and fold-back inversions (FBI). Under selection, genomic alterations coalesce into stable aneuploidy landscapes observed in almost all cancers [1– 7]. Previous research has mainly focused on measuring and interpreting aneuploidies, such as studies correlating copy number aberrations (CNAs) with markers of immune evasion and increased activity in proliferation pathways [7–11].

Research on the dynamics of CIN, whether chromosomal instability is actively ongoing at the time a tumour is sampled, has been less developed. A fundamental obstacle is that widely used copy number features, such as clonal diversity and subclonal heterogeneity, are insufficient to demonstrate ongoing CIN, as they can reflect historical events with subsequent clonal expansion [12, 13]; therefore, previous genomic studies have been limited to inferring CIN from these aggregate patterns [14–17] rather than directly measuring individual ongoing events.

Non-genomic approaches to measure ongoing CIN face inherent limitations in scale and resolution. Imaging-based approaches to observe mitotic errors require the processing of a large number of cells to capture the few that are undergoing mitosis[18, 19]. Three-dimensional live-cell imaging and single-cell karyotype sequencing of patient-derived monoclonal tumour organoid lines have been used to demonstrate the evolution of novel karyotypes over time *in vitro* [20, 21], but are limited in scale and genomic resolution and cannot be directly investigated in patient samples. Experimental systems and fluorescence in situ hybridisation (FISH)-based approaches offer controlled models or chromosome level insights [22–24], but neither provides a genome-wide measurement framework.

Recent advances in scWGS have opened the way to measuring CNAs in individual cancer cells at genome wide resolution [25–31]. Several methods have been developed for calling copy number profiles from scWGS data, including per-cell callers such as Aneufinder [32] and HMMcopy [33], and joint segmentation approaches such as CONET [34], SCICoNE [35] and SITKA [36]. Building on these tools, studies have leveraged scWGS to reveal subclonal diversity and distinct cell-to-cell mutational processes in cancer [37, 38], and shared CNAs have been used to infer evolutionary trajectories [39–41]. However, none of these approaches directly quantify the rate of ongoing CIN at the level of individual cells.

Here, we present and validate *scUnique*, a novel approach to measure ongoing CIN at scale by identifying individual recent copy number events from scWGS data. Our approach is conceptually similar to subclonality in bulk sequencing data [42], where the variant allele frequency of a CNAs indicates its relative time of appearance. For single cell data, we introduce the term recent copy number aberrations (rCNAs) to refer to CNAs that occur at low frequency within a cell population and have a greater probability of occurring temporally recently.

Measurement of ongoing CIN by rCNA requires overcoming three key challenges: (i) scWGS data are inherently noisy and sparse, requiring accurate per-cell copy number profiling; (ii) individual cells alone provide limited statistical power to localise breakpoints, necessitating information pooling across evolutionarily related cells; and (iii) candidate rCNAs must be rigorously distinguished from technical noise and phylogenetic inference errors. By overcoming these challenges, we establish a scalable, high-throughput method for measuring the dynamics of CIN via scWGS at high genomic resolution.

## Results

We give an overview of the *scUnique* approach and validate it (i) in a comprehensive benchmarking study, (ii) in CRISPR-Cas-based experimental systems, and (iii) single-cell derived clones of a well-characterised ovarian model. We then apply *scUnique* to distinguish ongoing HRD and FBI phenotypes.

### *scUnique* algorithm

The *scUnique* algorithm aims to identify recent CNAs (rCNAs) from a cell phylogeny by iteratively refining genome segmentation and tree reconstruction. This approach combines information across cells for highly accurate per-cell copy number profiles, given a phylogeny inferred from those profiles (Fig. 1). The approach can be divided into three separate phases: 1) initial per-cell segmentation; 2) refinement of per-cell segmentation using information from related cells; and 3) post-processing to identify only high-confidence rCNAs. We briefly describe the three phases and provide additional details in the methods section.

**Fig. 1:**
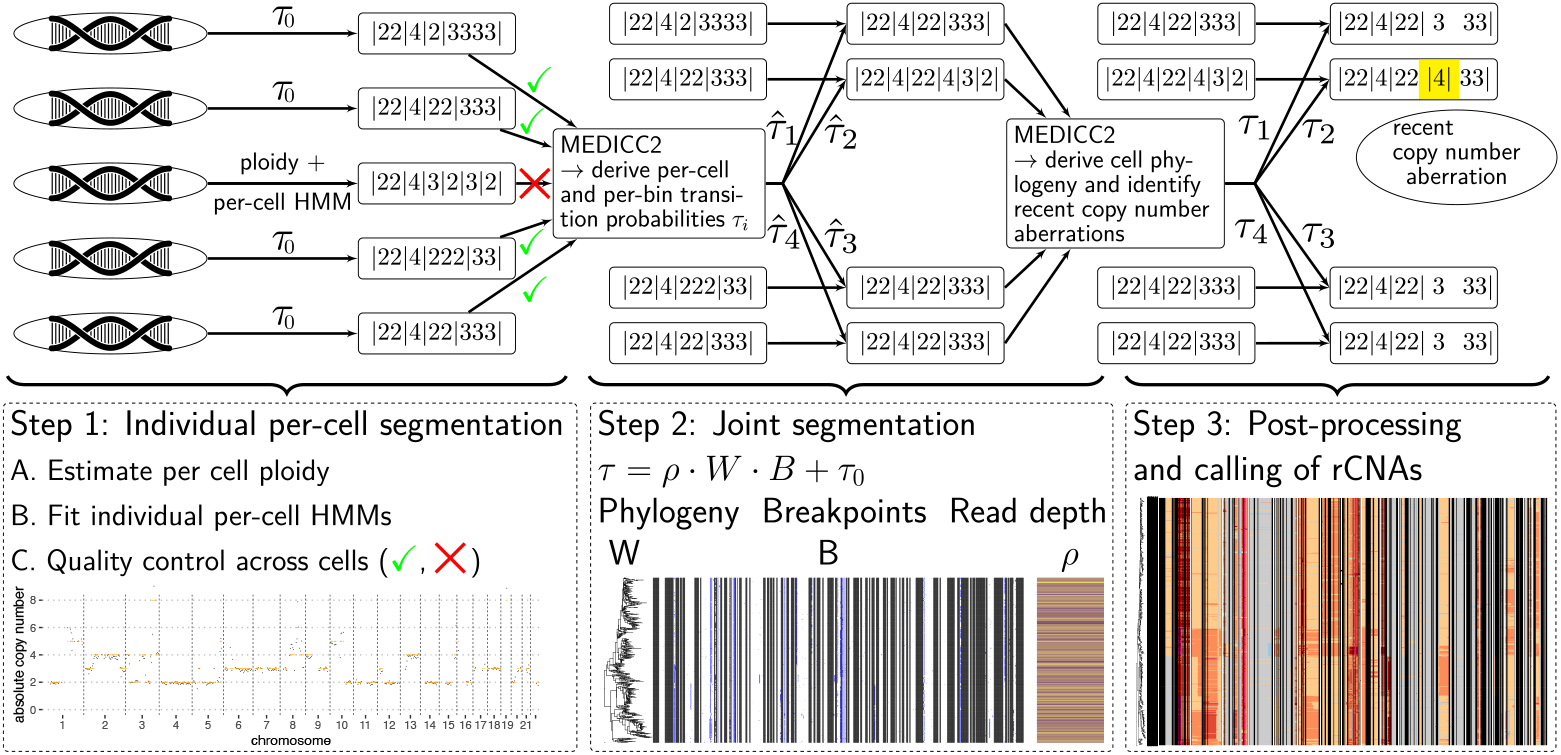
*scUnique* workflow. The method can be separated into three phases corresponding to the three panels at the bottom of the figure. Firstly, the cells are processed individually; per-cell ploidy is estimated, and a Hidden Markov Model (HMM) is fitted to each cell with a time-homogeneous and cell-invariant transition probability *τ*_0_ to determine a per-cell copy number profile. Replicating and other outlier cells are excluded from further joint processing. Secondly, information is pooled across related cells via updates to the per-cell copy number state transition probabilities *τ* based on information from related cells. Finally, using the phylogenetic relationship determined from the copy number profiles, cells that are closely related are scanned for copy number changes. We denote these changes as recent copy number aberrations (rCNAs).

Step 1. In the first step of *scUnique*, we use *scAbsolute* [43] for segmentation and ploidy estimation. *scAbsolute* begins with an initial segmentation of raw read counts per cell using a modified PELT algorithm with a negative binomial likelihood. This initial segmentation, referred to here as “scAbsolute-PELT”, facilitates the estimation of the read depth per-cell *ρ*_i_ in the next steps, which represents the scaling factor converting read counts to absolute copy numbers. Following this, *scAbsolute* includes an optional per-cell segmentation refinement step in which a HMM is fit. This HMM employs a position-invariant state transition matrix *T*_hom_, parameterised by the state transition probability *τ*_0_, and models the observed raw read counts as emissions from a negative binomial distribution conditioned on the underlying hidden copy number states. The refined segmentation obtained from this optional HMM step is referred to as “scAbsolute-HMM” here, which leverages the absolute copy number scaling inferred in the previous step. Notably, although the HMM segmentation relies on the ploidy estimate from the modified PELT segmentation, the two approaches differ fundamentally: PELT is a model-free changepoint detector that identifies breakpoints in raw count space, providing the segment structure required to estimate the scaling factor, whereas the HMM is a generative model that operates in the resulting absolute copy number space to assign integer copy number states with associated posterior probabilities.

Step 2. To refine segmentation by leveraging shared evolutionary information, we integrate data across all cells together. For this purpose, we run the MEDICC2 algorithm [41] to compute a pairwise distance matrix *W*, representing the phylogenetic relationship between cells. Using *W*, we update the per-cell, position-varying transition probabilities:

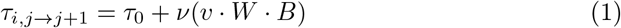

where *B* is a binary matrix of per-cell breakpoints from *scAbsolute, v* is a weight vector proportional to each cell’s read depth *ρ*, and ν is a normalisation function mapping values to the range [*τ*_0_, 1 − *ϵ*]. This yields an updated position-varying state transition matrix *T*_het_ (see Methods for details). This joint segmentation step yields more accurate copy number profiles than individual cell processing alone, as it leverages the phylogenetic structure to resolve ambiguous breakpoints and copy number states. Step 3. In a final step, we run MEDICC2 again on the jointly segmented cells to more accurately identify the phylogenetic relationship between cells. We use the final phylogenetic tree obtained at this stage to identify rCNAs. For each cell, we define candidate rCNAs as all copy number changes between that cell and its most recent ancestral node in the phylogeny, that is, changes that occurred on the terminal branch leading to that cell. We further filter candidate rCNAs to retain only those with high confidence (see Methods for details).

Applying MEDICC2 in the single-cell domain requires careful data preparation. Since MEDICC2 was designed for clonal copy number profiles, the input data must be of sufficient quality to ensure meaningful phylogenetic inference. To this end, we first remove cells undergoing DNA replication using the method described in [43]. We further apply stringent quality control to retain only cells with sufficient read depth and consistent normalised read coverage across the genome. Together, these filtering steps ensure that the copy number profiles used as input to MEDICC2 are reliable at the single-cell level.

### Validating *scUnique* on *in silico* data sets

To evaluate *scUnique* in comparison to existing copy number calling methods, we generated a ground-truth dataset from simulated copy number profiles (see Methods). In short, we layered individual, random, per cell copy number aberrations on top of simulated copy number profiles to investigate the potential for retrieving these simulated rCNAs. We systematically study the impact of read depth *ρ* (read depth normalised by bin size and sample ploidy) on the performance of copy number callers and how this impacts their ability to correctly identify copy number changes that occur at the individual cell level.

We use two different methods to simulate the copy number profiles to ensure diversity in the simulated genomes. Firstly, we simulate copy number profiles that are realistic but relatively homogeneous, comparable to a cell line setting or a single-cell derived clone. To do this, we performed single-cell whole genome sequencing on two ovarian adenocarcinoma cell lines (PEO1 and CIOV1) and generated a consensus copy number profile for each cell line using QDNAseq [44](see Fig. S1). Secondly, we use the simulator provided by the authors of SCICoNE [35] to create more complex and heterogeneous copy number profiles (SCICoNE 1-3). The corresponding simulation parameters are summarised in Table S1. We then simulate these copy numbers at different values of *ρ* (3, 5, 10, 15, 20, 30, 40, 50 and 100) to scrutinise the impact of read depth on the copy number callers.

For both simulation approaches (cell line-based and SCICoNE-based), we simulate individual, random per-cell copy number aberrations of varying size (2, 3, 5 and 10 bins) on top of the existing copy number profiles to investigate whether any of the methods are capable of retrieving these simulated rCNAs (see Figs. S2 and S3). Given the copy number profiles, we then draw the read counts from a negative binomial count model.

To evaluate the performance of *scUnique*, we compare against individual cell copy number callers: Aneufinder [32] and HMMcopy [33]. We also compare scUnique to our individual cell copy number calling methods used in Step 1 of the scUnique algorithm: scAbsolute-PELT [43] and scAbsolute-HMM. In addition, we compare against other joint copy number callers: CONET [34], SCICoNE [35] and SITKA [36].

We also evaluate three variants of *scUnique* that differ in how the cell distance matrix *W* is estimated in step 2: *scUnique*-UMAP (euclidean distance in UMAP space), *scUnique*-SCICoNE (phylogeny inferred by SCICoNE), and *scUnique*-MEDICC (MEDICC2 distance); see Methods for details.

When evaluating method performance, we focus on two aspects: overall accuracy in copy number calling, and detection of simulated rCNAs (see Fig. 2). Overall, *scUnique* performs best across the range of *ρ* values and across different simulations in terms of copy number accuracy (see Fig. 2a and Table S2). Across the methods tested, the accuracy of copy number calling generally increases with read depth, although there are some exceptions. For HMMcopy and methods that use HMMcopy for the initial ploidy estimation and segmentation (CONET and SITKA), the ploidy predictions vary based on the segmentation parameters used and the sequencing depth (*ρ*). This results in cases where incorrect ploidy predictions by HMMcopy contribute to lower overall performance at higher read depths when using the same segmentation parameters. SCICoNE, in the version tested here, contained a bug leading to worse performance for cells with higher read depth (author communication).

**Fig. 2:**
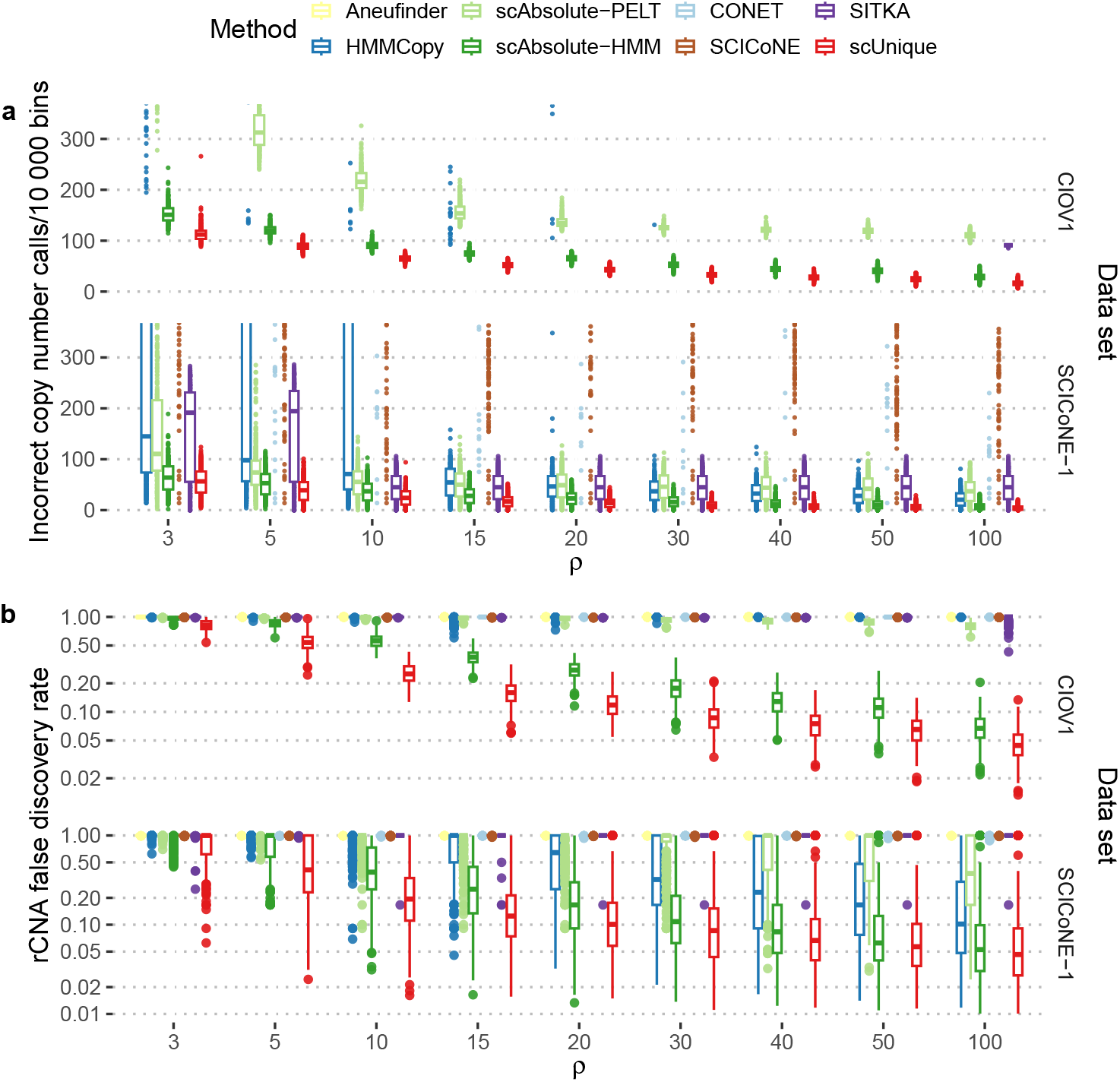
Method evaluation on simulated copy number profiles. Accuracy of copy number callers measured in two simulated data sets (CIOV1 and SCICoNE-1). Copy number callers for individual cells (Aneufinder, HMMcopy, scAbsolute-PELT, and *scAbsolute-HMM)*, and copy number callers using joint segmentation (CONET, SCICoNE, SITKA and *scUnique*) are assessed. **a)** The number of incorrectly classified copy number states per 10 000 bins. For clarity, only cells with less than 400 incorrect copy number calls per 10 000 are displayed here, for all results see Figs. S4 and S5. False discovery rate (FDR) for simulated rCNAs per cell for increasing read depth *ρ. scUnique* outperforms all other copy number callers in accuracy of copy number calling and in rCNA discovery in both simulated datasets. HMMcopy-based methods (HMMcopy, CONET, SITKA) see poor performance in the CIOV1 data set, both in terms of incorrect copy number calls and FDR which we determine is a result of calling an incorrect ploidy solution with the optimised set of segmentation parameters.

We find that the sensitivity of detecting rCNAs depends strongly on both read depth and event size (Figs. 2b and S6). For rCNAs covering only two bins, detection is very difficult even at high values of *ρ*, whereas larger rCNAs covering 10 bins can be detected at *ρ* values as low as 10. This dependence on event size directly informed the design of our simulation framework, in which we systematically varied the size of simulated rCNAs (2, 3, 5, and 10 bins) to evaluate method performance across a realistic range of event sizes. Across all event sizes, detection of rCNAs is poor at lower read depths (*ρ <* 30), and we observe FDR rates of 0.10 or lower rates that might be considered useful in practice for *ρ >* 30. Despite these challenges, *scUnique* and *scAbsolute-HMM* clearly perform best among the methods evaluated, with *scUnique* detecting larger rCNAs even at lower read depths where other methods fail (Fig. S6). We show that state-of-the-art performance can be achieved when information is integrated across cells, as demonstrated by the increased accuracy in copy number calling, and reduced FDR seen in *scUnique* (utilising jointly segmented copy number calling) over *scAbsolute-HMM* (per-cell copy number calling only) (see Figs. 2 and S7).

### Validating *scUnique* using CRISPR-Cas-based experimental systems

To validate the concept of rCNAs; that cell populations with high chromosomal instability can acquire sporadic, recent copy number aberrations, we apply *scUnique* to published single-cell DNA sequencing datasets from experimental models with impairments in DNA damage repair mechanisms. We apply *scUnique* to 184-hTERT mammary epithelial cells subjected to CRISPR-Cas9 editing to result in TP53 deficient (*TP53−/−*), TP53; BRCA1 deficient (*TP53−/−; BRCA1+/−* and *TP53−/−; BRCA1−/−*) and TP53; BRCA2 deficient (*TP53−/−; BRCA2+/−* and *TP53−/−; BRCA2−/−*) cell lines, published in Funnell et al. [38]. When comparing the number of rCNAs per cell, we see an increase in the number of events in *TP53* deficient cells compared to wild-type (KS test, *D* = 0.19, *p <* 10^−15^). In heterozygous knockouts of *BRCA1* and *BRCA2*, we see another small increase in the number of rCNAs per cell compared to TP53 deficient cells. (KS test, *D* = 0.19, *p <* 10^−15^). When comparing the number of rCNAs per cell in homozygous knockouts of *BRCA1* and *BRCA2* to their heterozygous counterparts, we see a marked increase in the number of rCNAs per cell (KS test: SA1188 (*TP53−/−; BRCA2+/−*) vs. SA1055 ((*TP53−/−; BRCA2−/−*), *D* = 0.32, *p <* 10^−15^, SA1292 (*TP53−/−; BRCA1+/−*) vs. SA1054 (*TP53−/−; BRCA2−/−*), *D* = 0.68, *p <* 10^−15^. Similar to the number of rCNAs per cell, we observe an increase in the relative number of cells that harbour any rCNA, with 99.6% of cells that are homozygous BRCA1 knockout (SA1054) having some rCNAs. In comparison, less than 15% of wild-type cells have any rCNA, as seen in the TP53-proficient fibroblast (3.9%) and the SA039U epithelial wild-type cells (12.7%) (Fig. 3e). Note that we also observe a difference in the number of rCNAs between the two latter types of “normal” cells (KS test, SA039U vs. Fibroblast, *D* = 0.08, *p <* 10^−7^), indicating cell-type specific base rates of mutational activity.

**Fig. 3:**
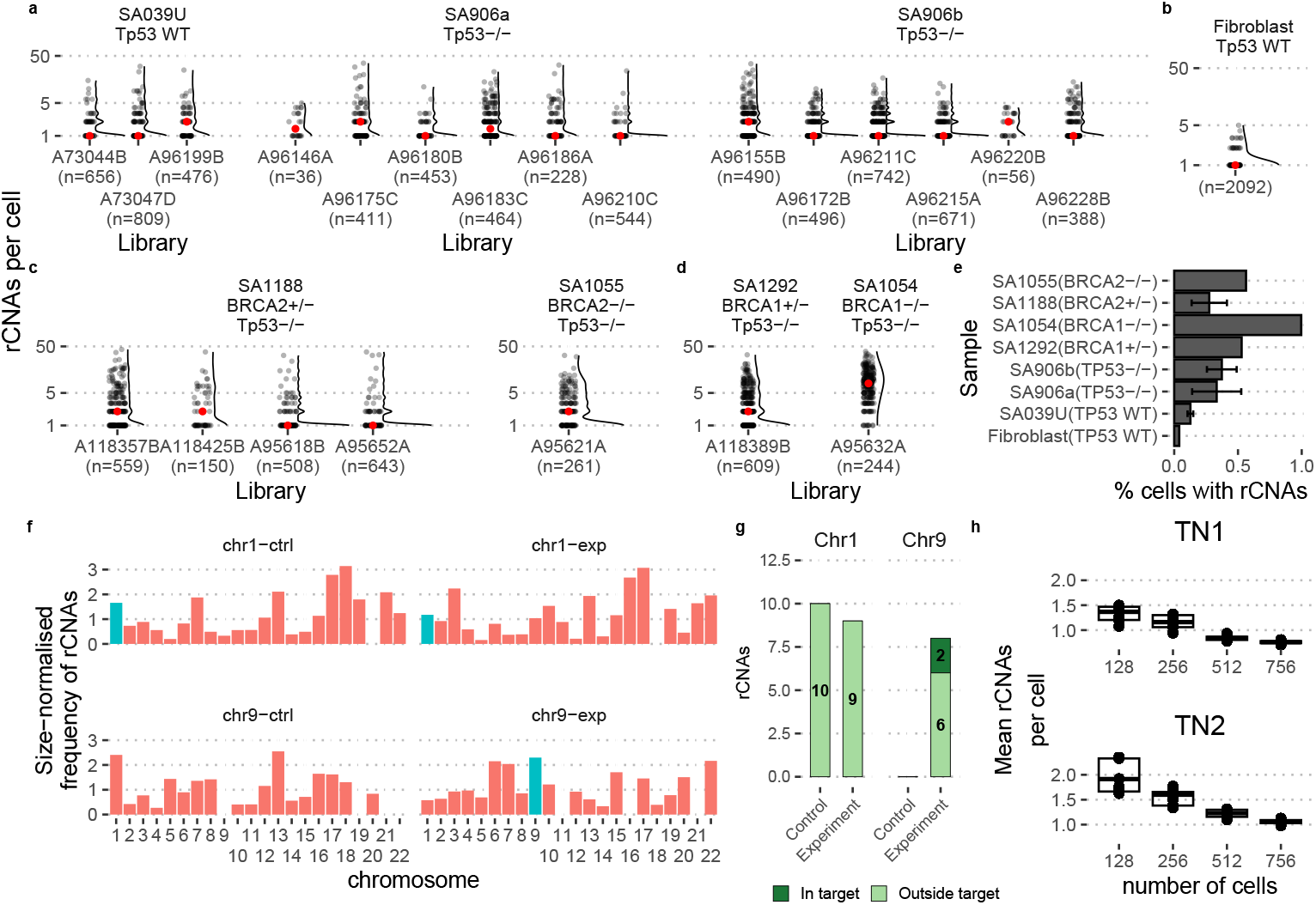
rCNAs in experimental systems with CRISPR-Cas induced chromosomal instability. **a-d)** Number of rCNAs in TP53 wild type (SA039U, Fibroblast), TP53-deficient cells (SA906a/b), with additional heterozygous (SA1188, SA1292) and homozygous (SA1054, SA1055) knockouts of BRCA1/BRCA2 genes. The red point indicates the median of the discrete distribution on the left, that is approximated by the violin plot on the right. **e)** The proportion of cells harbouring at least one rCNA for the different data sets from **a-d.**Error bars indicate ± one standard deviation when multiple libraries are available for a sample. **f)** Size-normalised frequency of rCNAs in experimental system to induce increased CIN using nuclease-dead CRISPR-Cas9 targeting specific genomic loci (Chr 1, Chr 9). Frequency of rCNAs is increased on chromosome 9 compared with the control (Chr9-exp vs Chr9-ctrl), but no such effect is visible for the experimental manipulation of chromosome 1 (Chr1-exp vs Chr1-ctrl). Experimentally manipulated chromosomes (Chr1, Chr9) are marked in cyan. **g)** Distribution of rCNAs on targeted chromosomes split by location: within the CRISPR-targeted region (dark green) versus elsewhere on the chromosome (light green). CRISPR-targeted samples show enrichment of events within the targeted regions compared to controls. **h)** TN1 and TN2 tumour samples with originally 1100 and 1024 cells, respectively, are randomly subsampled ten times each to 128, 256, 512, and 756 cells. The average number of rCNAs decreases with increasing sample size.

We next examine whether our approach can reliably detect experimentally induced CIN in specific genomic locations. Tovini et al. [22] have developed a method to induce targeted aneuploidies by fusing nuclease-dead CRISPR-Cas9 (dCas9) with the kinetochore-nucleating domain CENP-T, targeting either pericentromeric repeats on chromosome 9 (Chr9-CEN) or telomeric-proximal repeats on chromosome 1p (Chr1-TELO) in HEK293T, a human embryonic cell line with a heterogenous, hypotriploid genomic background.

We validate our method by specifically looking at the relative frequency of rCNAs on these two chromosomes (see Fig. 3f). Despite the modest number of cells (32 cells per condition), we find clear evidence of ongoing CIN on chromosome 9 as a consequence of the experimental manipulation (*z* = − 2.258, *p* = 0.012; 8 vs 0 events in control and experiment, respectively) but no increase in events located on chromosome 1 (*z* = − 0.800, *p* = 0.788; 10 vs 9 predicted rCNAs in control and experimental conditions, respectively)

Only 2/8 Chr9 (Fig. 3g) rCNAs localise directly to the targeted pericentromeric region, with the remaining events distributed along the chromosome. This pattern is consistent with ectopic kinetochore–induced chromosome bridges seeding secondary rearrangements, as observed for bridges that initiate breakage–fusion–bridge cycles and additional segmental events in other systems [45], rather than exclusively producing focal lesions at the targeting site. In contrast, no Chr1 rCNAs (0/10 experiment, 0/9 control; Fig. 3g) coincide with the telomeric target, reflecting a combination of heterogeneous distant breaks, weaker KNL1 recruitment (3x on Chr1 vs 20x on Chr9 compared to endogenous levels), and Chr1’s high baseline subclonality that saturates predictions.

Unlike the >20 Mb threshold used by Tovini et al. [22] which misses finer CIN, our method resolves distributed recent events across Chr9, confirming ongoing instability while Chr1’s null aligns with their reported noise and modest effects.

A limitation of our approach is that the frequency of rCNAs is only a relative measure of time and partially depends on the size of the sample. In order to understand how strongly the number of rCNAs is impacted by the number of cells in a sample, and how robust our measure is to the randomness inherent to sampling, we conduct a downsampling experiment. We use two triple-negative breast cancer (TNBC) tumours from the MD Anderson Breast Tissue Bank: TN1 (chemotherapy-treated Ductal Carcinoma In Situ (DCIS); 1100 cells) and TN2 (untreated Invasive Ductal Carcinoma (IDC); 1024 cells), sequenced via Acoustic Cell Tagmentation (ACT), a high-throughput, single-molecule scWGS method [46]. We subsample 10 random sets consisting of 128, 256, 512 and 756 cells each and run *scUnique* on the subsets. Fig. 3h shows that the estimate of rCNAs per cell is relatively robust between random samples and across different sample sizes. Increasing sample size leads to an overall increase in the number of rCNAs, while reducing the number of rCNAs observed per cell. In the hypothetical limit of an infinite sample size and a perfect phylogeny, we would expect to observe all CNAs that are a result of the latest cell division.

### *scUnique* discovers ongoing as opposed to historical markers of chromosomal instability

Following validation on *in silico* datasets and CRISPR-Cas9 based experimental systems, we investigated whether *scUnique* identifies *ongoing* chromosomal instability rather than historical events inherited from ancestral cells. To test this, we cultured six single-cell-derived clones (B4, C9, D5, D7, D11, G3) from a common PEO1 parent sample and performed scWGS to measure rCNAs arising in each clone (Fig. 4a). We performed scUnique on a subset of 15 cells per clone (90 cells total) and 15 cells from the parent population.

**Fig. 4:**
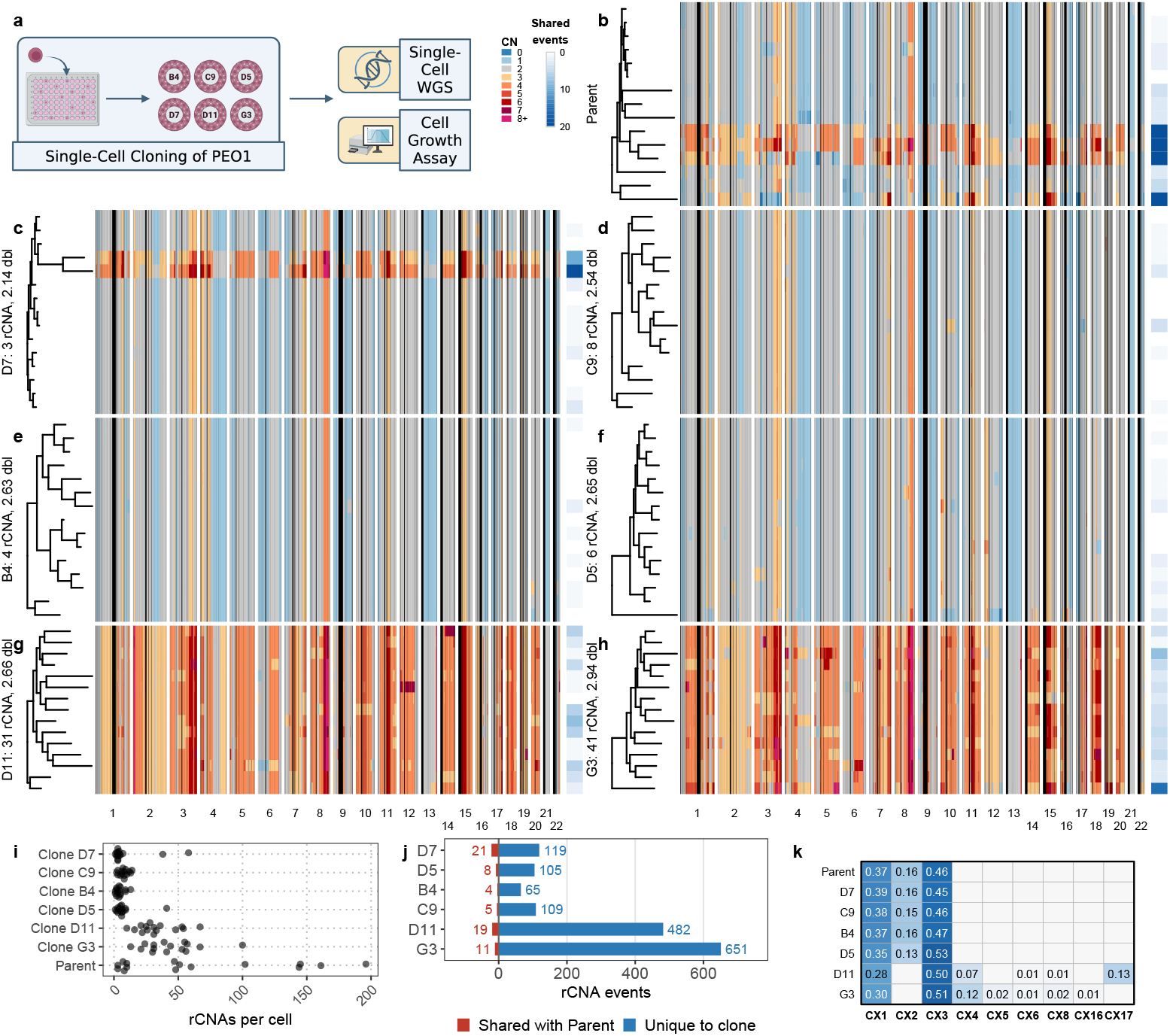
Validation of *scUnique* rCNA detection using clonal proliferation assays. **(a)** Experimental workflow: six single-cell-derived clones (B4, C9, D5, D7, D11, G3; 15 cells each) were expanded from a common PEO1 parental cell line (15 cells). Cell doublings were calculated from longitudinal confluence measurements, and single-cell DNA sequencing was performed. Figure created using BioRender. **(b)** Copy number heatmap of the parental PEO1 population showing baseline genomic heterogeneity. The right annotation bar indicates, per cell, the number of rCNAs that are also detected in at least one clone (shared events; white–blue gradient, 0–20). **(c–h)** Copy-number heatmaps for each clone ordered by proliferation rate from slowest (D7, 2.14 doublings) to fastest (G3, 2.94 doublings). Rows represent individual cells clustered by MEDICC2 phylogenetic trees; columns represent genomic bins. Clone labels indicate median rCNA count and doubling time. The right annotation bar indicates the number of rCNAs per cell that are shared with the parental population (white–blue gradient, 0–20). **(i)** Distribution of rCNAs per cell across parent and clones. Points represent individual cells. **(j)** Clone-level decomposition of rCNAs: events shared with the parent (left) versus events unique to each clone (right). **(k)** Heatmap of pseudobulk chromosomal instability signature (CX) activities across parent and clones.

rCNA burden differed markedly across clones. D11 and G3 showed the highest per-cell burdens (Fig. 4i), consistent with their highly disordered copy-number heatmaps and whole genome doubling (WGD)-associated instability (Figs. 4g, 4h and 4k). D7 showed a low median burden overall (4 events/cell) but contained two outlier cells with high counts (38 and 58 events; Fig. 4c). These outliers had catastrophic features: 27-fold larger events than baseline (median 13.6 Mb vs 0.5 Mb) and increased centromere-proximal enrichment (17.7% vs 9.1% within 5 Mb), suggesting mitotic defects or centromere dysfunction as triggering mechanisms.

We then tested whether proliferation rate could explain these differences. Cell doublings were calculated from Cell Titer Glo measurements (log_2_(T_7_*/*T_0_), where T_0_ = 24h and T_7_ = 168h), ranging from 2.14 (D7) to 2.94 (G3), despite shared ancestry (Table S3). Median rCNA burden showed a positive clone-level association with doublings (Spearman *ρ* = 0.829, *p* = 0.0583; Fig. S8a), but this association attenuated in ploidy-adjusted multivariable model, with ploidy remaining significant.

Event-sharing analysis provided direct evidence for recent acquisition (Figs. 4j and S8b). Clone-to-clone sharing was minimal (mean 1.3%), and only 48 of 1,542 clone events (3.1%) appeared in multiple clones, supporting largely independent evolution after clonal isolation. Shared recurrent events mapped preferentially to chromosomes 3, 9, and 17, consistent with fragile genomic hotspots. Clone-to-parent sharing was modestly higher overall (mean 6.3%) and was enriched in WGD-bearing entities: the two D7 outlier cells and WGD-dominant clones D11 and G3 showed elevated clone-to-parent sharing (20 and 10 events for D7 outliers; 19 and 11 for D11 and G3, respectively; Figs. 4c and S8b).

To characterize the clones and their differences, we aggregated cells within each clone into pseudobulk copy-number profiles and applied copy-number signature analysis [17]. Together, CX4 signature activity in D11 and G3 (0.075 and 0.117, respectively; Fig. 4k) and Fluorescence-activated Cell Sorting (FACS)-confirmed polyploidy (Fig. S15) in the parental population support an ongoing WGD process represented across successive clonal states: unaffected diploid populations (B4, C9, D5), early-stage involvement (D7, 2/15 cells affected), and clonal dominance (D11 and G3, 15/15 cells affected). This stage-structured heterogeneity is difficult to reconcile with a single fixed historical event. Consistently, pseudobulk copy-number signatures were highly similar across the four diploid/transitional clones (B4, C9, D5 and D7), indicating a shared mutational-process background, whereas the WGD-dominant clones showed additional activity of signatures associated with post-WGD genomes (D11: CX3, CX6, CX8, CX17; G3: CX3, CX5, CX6, CX8), with no CX2 activity in either clone. Critically, *scUnique* detected a high rCNA burden in the D7 outlier cells that was not captured by the pseudobulk signatures, highlighting the added resolution of single-cell analysis.

To confirm that rCNAs reflect genuine instability rather than artifacts of phylogenetic inference, we compared rCNA burden with orthogonal instability metrics computed directly from raw copy-number profiles. Across cells from all six clones, rCNA burden correlated strongly with copy-number variance (*ρ* = 0.78, *p <* 0.001), Shannon entropy (*ρ* = 0.76, *p <* 0.001), and breakpoint count (*ρ* = 0.74, *p <* 0.001; Fig. S9). Collectively, these results indicate that *scUnique* captures ongoing chromosomal instability during clonal evolution rather than merely cataloguing historical clone-defining CNAs.

To assess whether rCNAs reflect ongoing instability rather than artifacts of phylogenetic inference, we compared per-cell rCNA burden with orthogonal instability metrics derived directly from raw copy-number profiles. Across pooled cells from all six clones, rCNA burden was positively correlated with copy-number variance (*ρ* = 0.78, *p <* 0.001), Shannon entropy (*ρ* = 0.76, *p <* 0.001), and breakpoint count (*ρ* = 0.74, *p <* 0.001; Fig. S9). However, these associations were heterogeneous across clones and were strongest in the most copy-number-disordered clones, indicating that the pooled correlations are driven in part by between-clone differences. Thus, *scUnique* appears to capture biologically meaningful variation in chromosomal instability, although the strength of this relationship is context dependent rather than uniform across clones.

### Mutational processes of chromosomal instability through the lens of rCNAs

We then assessed whether ongoing CIN, as defined by rCNAs, reflected distinct mutational processes. We begin to address this using a published cohort of samples with known and relatively distinct mutational signature activity [38]. The data set comprises 28 Direct Library Preparation (DLP)+ samples with sufficient cell numbers and read depth for our analysis: 23 high-grade serous ovarian carcinoma (HGSC) and 3 TNBC cases from patient-derived xenografts (PDXs) and patient tissues, and two 184-hTERT mammary epithelial cell lines with CRISPR-Cas9 mediated *TP53−/−* and *BRCA1−/−* knockouts. Samples are characterised by HRD or FBI phenotypes (16 and 12 samples, respectively) as determined through bulk level analysis of mutational processes. One major limitation of this data set is the confounding of ploidy with mutational signature status. Most of the high ploidy samples are characterised by FBI, whereas the low ploidy samples are exclusively characterised by HRD signature (Fig. 5d).

**Fig. 5:**
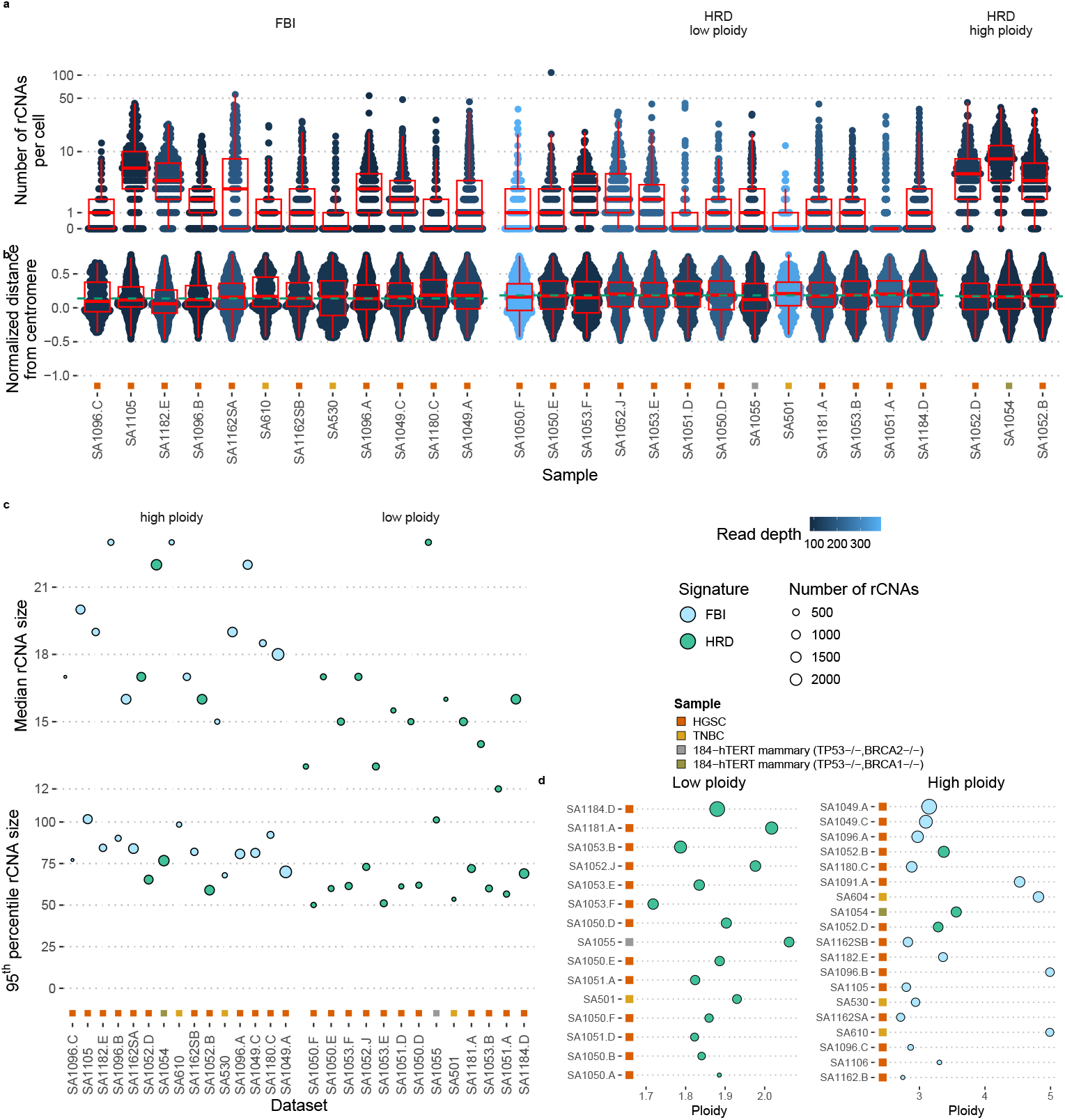
rCNA-based features differ between samples with foldback inversion (FBI) and homologous recombination deficiency (HRD) signatures of mutational processes. Samples characterised by bulk-sequencing derived mutational signatures [38] of mutational processes (HRD and FBI signatures). **a)** Each point indicates the number of rCNAs per cell. **b)** Each point indicates the relative (normalised by chromosome size) distance of an rCNA from the centromere region. Positive values indicate position of the rCNA on the larger q arm, and negative values indicate the rCNA is on the p arm. Dashed green lines indicate mean normalised distance per group (FBI, HRD low ploidy, HRD high ploidy). **c)** Size of rCNAs in MB. Each point indicates a data set, with the number of cells denoted by the size of the radius of the circle, and the colour indicating HRD vs FBI phenotype. **d)** Sample ploidy for the different data sets included in the study. Note the difference in ploidy between the FBI and HRD signature samples. We use a cut-off of ploidy 2.5 to distinguish between low and high ploidy samples. Samples with less than 75 cells harbouring rCNAs are not included in the analysis and in **a-c)**.Samples are sorted by increasing number of rCNAs within each group.

We observe a varying number of rCNAs per cell (Fig. 5a), with no clear difference between the two groups (Welch’s t-test, *t* = − 0.295, *p* = 0.769). However, the two groups differ in the distance of the rCNA from the centromere (see Fig. 5b, Welch’s t-test, *t* = − 4.07, *p* = 2 × 10^−4^). Note that we normalise the distance by chromosome size to avoid any bias arising from different chromosome sizes.

We also see a difference in the size distribution of rCNAs (see Fig. 5c), with rCNAs in the case of HRD characterised samples tend to be smaller, while FBI samples show a tendency for larger size rCNAs as measured by the median size of a sample (Wilcoxon rank sum test, *W* = 367, *p* = 0.003) and by 95-percentile of a sample (Wilcoxon rank sum test, *W* = 439, *p* = 3.96 × 10^−6^). The observed median rCNA size ranges from about 12 − 18 Mb across the HRD associated samples, which is similar to the size of genomic segments characteristic of mutational processes associated with HRD (segment sizes ranging from 2.5 Mb to 30 Mb for HRD associated signature CX3, as described in Drews et al. [47]).

## Discussion

*scUnique* offers a novel perspective on CIN by characterising CNAs at the level of individual cells. By integrating copy number information across cells and inferring phylogenetic relationships based on CNAs, we show that we can accurately detect rCNAs in scWGS data. We demonstrate that rCNAs are a marker of ongoing CIN and can be used to quantitatively measure the dynamics of CIN across cell lines and tumour samples.

Measurement of ongoing CIN at the time of sample collection has major implications for therapy decisions. Mutational signatures of chromosomal instability have shown exciting promise as new genomic biomarkers for the underlying mechanisms resulting in CIN; one example is scoring HRD and subsequent treatment with PARP inhibitors [47–50]. However, their use of bulk sequencing methods is a limitation; only mutational processes present in high-frequency cell populations are detected, with rarer, low-frequency events not detected. This limits the detection to processes that have undergone clonal expansion, and therefore represent a historic view of the tumour. This severely limits their ability to detect recent changes in the activity of mutational processes. The activity of mutational processes has been shown to vary over time [20, 51]; they can be inactivated, for instance via a reverter mutation in BRCA1 or BRCA2 restoring homologous recombination repair mechanisms [52–55]. Importantly, historical markers of mutational signatures, such as CNAs or single-nucleotide variations (SNVs) can be present in a sample in the absence of the active mutational process which could lead to an incorrect treatment decision [56].

What is lacking is a quantitative whole-genome readout of current CIN activity. While Hernando et al. recently proposed an approach for finding cell-unique events in scWGS data [57], their unpublished methodology precludes direct comparison. rCNAs have the potential to act as dynamic biomarkers for the presence of active mutational processes. This notion is particularly relevant in cases of relapse or resistance, where we would expect changes in biological mechanisms to appear over a background of historical genomic changes.

The concept of rCNAs can improve our understanding of the biological mechanisms and generative process underlying chromosomal instability. Current approaches are limited in terms of temporal resolution, and it is consequently difficult to separate effects of fitness, and interaction between different biological mechanisms. By measuring CIN at scale, at a whole-genome resolution, and with high temporal resolution, we can learn how individual CNAs contribute to the genomic complexity that we observe at the bulk sequencing level. We postulate that rCNAs are less impacted by fitness effects and interaction between different CNAs because relatively little time has passed since their acquisition. Observing the elementary changes at the single cell level should therefore help unravel the contribution of the individual factors more clearly. However, further work will be required to characterise the contribution of individual mutational processes to the patterns of rCNAs observed.

One important practical aspect of this study is the systematic investigation of read depth required to obtain reliable copy number estimates in single-cell DNA sequencing data. We provide evidence guiding the design of future sequencing studies: either in terms of sequencing depth required to obtain accurate copy number inference given a genomic bin size, or equivalently the genomic resolution that accurate copy number inference is possible at, given a certain sequencing depth.

The notion of cellular copy number variation in the form of rCNAs is an extension of the concept of clonal and subclonal copy number variation to the single cell limit. While we compare a number of methods in this study, it is important to note that the current method is to our knowledge the first method explicitly aiming to uncover genomic changes at the level of individual cells. While previous work has demonstrated the existence of cell-to-cell variation at the level of structural and copy number alteration [38], here we provide a tool to systematically study this variation, taking into account an evolutionary model of copy number changes.

None of the joint copy number calling methods tested here were developed for the purpose of identifying rCNAs; therefore, the low performance in this particular task is not surprising. Interestingly, when HMMcopy identifies the correct ploidy, it also performs well and outperforms the joint copy number callers in terms of rCNA detection. This possibly indicates that HMM-based methods are generally well suited for this problem, and that integration approaches developed so far were not optimized for sensitivity to low-frequency events.

There exist a number of limitations for our approach:

First, *scUnique* can not distinguish between individual CNAs and a single CNA comprising multiple genomic loci. The method is naturally limited by the choice of genomic bin size and larger CNAs are generally much easier to detect, and are therefore likely to be over-represented. Obtaining higher genomic resolution is possible, but comes at the expense of deeper sequencing and higher cost.

Second, we cannot give an absolute time resolution for individual rCNAs. Instead we rely on the relative notion of recency. Further work will be required to get a better understanding of the time scales and relationship between cell doubling times and the rates at which rCNAs occur. We have demonstrated that our method is robust to variation in the number of cells, but the temporal resolution naturally improves with the number of cells in a sample.

Third, we do not estimate allelic copy numbers in this study. While this would be useful in improving the quality of the phylogenetic tree and helpful in identifying CNAs occurring under parallel evolution, we chose the more generic approach that is applicable to all our data sets.

Fourth, the model could be further improved using prior information to choose the value of *τ*_0_, or by introducing copy number state-specific transition probabilities. We decided against including prior information in this work, in order not to introduce potential biases stemming from bulk as opposed to single-cell analysis, and due to the lack of deeply sequenced scWGS data sets that would allow us to estimate these parameters.

Finally, it is important to note that using our method we are only able to make statements about an ensemble of rCNAs. We are able to make statistical statements and compare subpopulations of cells, however, we do not make claims about individual CNAs in individual cells. Only in observing many cells and CNAs, we can reasonably control the false discovery rate and ignore individual errors in placing cells in the phylogenetic tree.

Despite these limitations, *scUnique* represents a significant advance in measuring the dynamics of CIN at the single-cell level, providing a scalable, genome-wide framework to distinguish ongoing chromosomal instability from historical events. By enabling quantitative measurement of rCNAs, *scUnique* lays the groundwork for the development of dynamic biomarkers with the potential to inform treatment decisions in the future.

## Methods

### *scUnique* algorithm

In the following, we provide a detailed description of the three phases of the *scUnique* algorithm, starting with the statistical model and per-cell processing, followed by the joint segmentation refinement, and concluding with the identification of rCNAs.

### Setting

We first define the notation and the read count model used throughout. We split the genome into *M* fixed-size genomic bins. We refer to the (unknown) per-bin copy number as *c*_i,j_, its estimate as ĉ_i,j_, and the read count observed in bin *j* in cell *i* as *x*_i,j_ (*j* ∈ [1, *M*], *i* ∈ [1, *N*]). We model the read count distribution with a negative binomial distribution [58] (see Fig. S10 for a rationale):

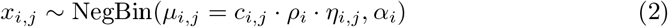

We parametrise the negative binomial distribution via its mean *µ* and an overdispersion parameter *α*. The overdispersion parameter characterises the relationship between mean *µ* and variance of a random variable X that follows a negative binomial distribution: Var(*X*) = *µ* + *α* · *µ*^2^. Hence, for values of *α* → 0, the distribution approximates the Poisson distribution.

We model the mean observed read counts as a linear function of copy number state *c*_i,j_ of a given bin, the read depth of the cell via *ρ*_i_ and a multiplicative GC and mappability correction factor *η*_i,j_ centred around 1. *ρ*_i_ denotes the average read depth per copy and per bin for cell *i. ρ* is a measure of per-cell read coverage that we find very useful to compare different cells in terms of read depth, independently of cell ploidy and copy number profiles. The value of *ρ* is a direct measure of the expected difference in mean read counts between neighbouring copy number states. The relationship between GC content and mappability, and observed read count is modelled individually per cell. Similarly, we assume the overdispersion parameter *α* varies across cells. In practice, *α* appears to depend both on the sample and the sequencing technology used, but its variation is relatively small and it constitutes a technological limitation (Fig. S11), unlike the sequencing depth represented by *ρ*, which can be adjusted as part of the experimental setup. Consequently, we focus our analysis on the impact of read depth *ρ*.

### Per-cell ploidy values

We make use of a reliable estimate of *ρ*, referred to below as 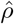, that is obtained via running scAbsolute [43]. The fundamental advantage of running the inference algorithm with a well-initialised and constrained value of *ρ* lies in avoiding the symmetries in the likelihoods that exist when the ploidy is not known. As a simple example, consider the case of a normal, diploid genome. For an HMM, the solution in terms of different copy number states is equivalent, in terms of the likelihood, as long as *ρ* is not known, e.g. there is no way to distinguish a diploid from a tetraploid genome. This corresponds to a case of 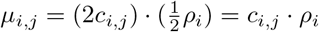, for a value of *c*_i,j_ = 2. In other words, the above model is ill defined because of multiplicative factors.

We completely avoid this problem with our approach. In addition, there is no dependency between the segmentation parameters and the predicted ploidy solution, as observed in other methods (see Figs. 2a and S4). As part of the ploidy estimation procedure, we obtain an initial segmentation relying on the PELT algorithm for change point detection [59]. Our PELT implementation uses a simple Method-of-Moments estimator for the parameters of the negative binomial likelihood [60, 61].

We found this estimator to be useful in practice, as it provides a good trade-off between speed and accuracy, and estimates the segment means rather than copy number state. *scAbsolute-PELT* is included as a baseline scenario in the method evaluation and corresponds to the segmentation obtained in Schneider et al. [43].

### Per-cell GC and mappability correction

It is well known that GC content and mappability variation across genomic bins have a marked impact on the observed read counts [28, 44]. To obtain more accurate copy number predictions, we estimate a correction factor to estimate how these factors impact observed read counts similar to the cell cycle correction described in Schneider et al. [43]. We model the relationship with a generalised additive model (GAM) in order to obtain a GC and mappability correction factor *η*_i,j_ separately for each cell and bin.

We use the PELT segmentation to account for differences in underlying copy number state when estimating the GC and mappability correction. The following describes the model used for the counts in a given cell *i* at bin *j*:

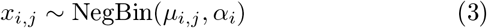

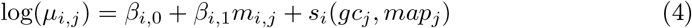

where *m*_i,j_ refers to the mean count levels for a segment covering bin *j* obtained via the PELT segmentation, and *s*_i_(*gc, map*) describes the smooth functions used to model the impact of GC content and mappability in cell *i*. We use smooth plate thin splines for the function *s*. Our modelling approach does not require the segmentation to be exact, as long as it is correct in the majority of cases due to the smooth modelling. By including the segmentation as a covariate in the model, we account for the fact that while a significant part of the variance within a given segment can be explained by GC and mappability variation, the variation between segments is dominated by copy number state. The underlying assumption is that the effect that GC and mappability variation have on the observed read counts is small compared to a change in copy number state. This model directly links the mean of the observed read counts with the effect of GC content and mappability via a multiplicative factor *η*_i,j_ = *s*_i_(*gc*_j_, *map*_j_). The factor is estimated individually per cell, and varies across bins on the basis of their GC content and mappability.

### Individual per-cell segmentation

In the *scUnique* approach, we initially fit an HMM separately to all cells, disregarding the information from other cells. We initialise *ρ*_i_ with our estimate 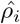 obtained through the ploidy estimation procedure described in [43]. Similarly, we precompute the GC and mappability correction factor *η*_i,j_ as described above using the PELT segmentation.

While this approach means that the useful information from other cells is not available initially, it has several advantages: i) The initial computational step is trivial to parallelise. ii) Prior to any downstream joint analysis, we can exclude cells that are undergoing DNA replication and/or are of low quality – to reliably obtain this information requires some preprocessing and segmentation. iii) We avoid any pre-selection of breakpoints, that is often used to reduce computation time.

We specify the following model for the HMM. The underlying hidden states *c*_i,j_ in cell *i* at position *j* correspond to the copy number states, running from 0 to a maximum cutoff copy number state *c*_max_. We choose a uniform initial state distribution. We initialise the time-varying emission probabilities as

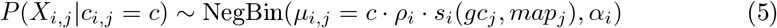

The correction factor *s*_i_(*gc*_j_, *map*_j_) = *η*_i,j_ for the influence of GC content and mappability variation is fixed at this stage. In the case of copy number state *c* = 0, we estimate a separate set of parameters: 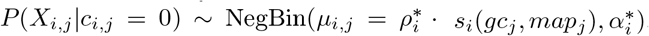.

Overall, we estimate only four parameters per cell: *ρ, α* and a separate *ρ*^*^ and *α*^*^ for the zero-copies state. Crucially, because we have already obtained a good enough fit to determine the correct ploidy solution up to a small error, we can initialise the HMM parameter *ρ* with a very good initial estimate 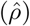, avoiding the problem of unidentifiability.

The remaining issue that any segmentation algorithm has to address is the trade-off between sensitivity and specificity, or in other words: how many segments to call? In the case of HMMs, we make this decision via the state transition matrix *T*. Choosing an optimal threshold is an open research question. In the single-cell segmentation case, we choose to parametrise the copy number state transition matrix *T* with a single parameter *τ*_0_ for all values of *τ*_i,j→j+1_, and use a homogeneous time-transition, i.e. the value of the transition matrix *T* = *T*_hom_ is identical along the length of the genome, independent of bin position *j*.

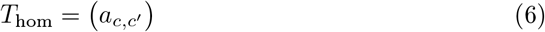

where

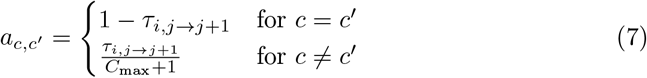

In the case of *T*_hom_, we choose *τ*_i,j→j+1_ = *τ*_0_ ∀*j*. By varying the value of *τ*_0_, we tend to create more segments (*τ*_0_ → 1), or we can enforce a more conservative segmentation (*τ*_0_ → 0). We purposefully make a transition from a given state to any other state equally likely.

In practice, throughout we choose a constant value of *τ*_0_ = 10^−3^, to avoid oversegmentation and only accept copy number changes based on strong evidence. This initial value is chosen to provide a conservative (not necessarily optimal) segmentation and we find that this value performs reasonably well. We did not optimise the value of *τ*_0_ as part of our simulations, because of the additional computational cost this would incur.

### Quality control

#### Bin-level quality control

We rely on a selection of high-quality bins described in [43]. The notion of selecting high quality bins is to assure the linear relationship between read depth and copy number state that our method relies on. The selection of bins is mostly based on public single-cell sequencing data of normal, diploid cells with DLP and 10x sequencing protocols and is used consistently throughout all analyses. We exclude the bins on the X and Y chromosome from the joint analysis, as we find the read mapping on these chromosomes generally less reliable.

#### Removal of replicating cells

Prior to discarding any outlier cells, we remove cells undergoing DNA replication based on the method described in [43]. This is particularly relevant in cell line data, where a large proportion of cells is undergoing DNA replication at any time point. Cells undergoing DNA replication tend to harbour a large number of ephemeral copy number aberrations that occur as part of the DNA replication process, and could be confounded with rCNAs.

#### Cell-level quality control

Quality of single-cell sequencing libraries is varying and one commonly observes a number of low quality cells. However, it is relatively difficult to exclude cells based on a single quality measure [32]. While it is possible to train a classifier to automate quality control [28], this requires a significant time-investment in labelling cells. Furthermore, in the author’s own experience, manual detection of outlier cells based on visual inspection of copy number profiles can be surprisingly difficult, as it is highly dependent on read coverage and copy number profile. Finally, it is entirely unclear how well such classifiers generalise to novel data sets and different sequencing technologies.

As a consequence, we opt for an alternative approach using two common quality metrics - MAPD and Gini coefficient - to identify outlier cells.

Both metrics measure how evenly reads are distributed across the genome, and have been previously used for the purpose of cell-level quality control [62, 63]. We explicitly normalise the two metrics to account for the effect of read depth; in the case of Gini coefficient we also try to normalise to account for differences in copy number profiles. By normalising these values, we aim to overcome some limitations of both measures when dealing with heterogeneous samples (see Fig. S12).

The MAPD (Median Absolute deviation of Pairwise Differences) statistic [62] is defined as:

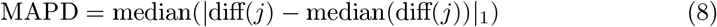

where 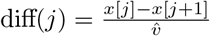, where 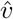 is defined as the mean of the vector *x*. While the MAPD statistic is relatively robust to differences in copy number profiles, its value is dependent on the cell’s read depth (see Fig. S12).

Secondly, we compute the Gini coefficient detailing the distribution of reads across the genome. A Gini coefficient of 0 expresses perfect equality, where all values are the same. A Gini coefficient of 1 expresses maximal inequality among values. The distribution of reads in this setting depends on the copy number profiles of the cells observed. The Gini coefficient shows strong variation based on the underlying copy number profile. This makes it unsuitable for analysing heterogeneous samples. In order to account for this variation, we introduce a normalised Gini coefficient. The normalized per-cell Gini coefficient is created by computing a Gini coefficient per segment and then computing the segment-length weighted mean across all segments.

For both metrics (MAPD, Gini), we use a robust linear regression model to regress out the influence of read depth *ρ* on the measure. We use the standardised residual to identify and remove outlier cells (by default with a cutoff value of 1.5).

A cell is considered an outlier, if it is identified as an outlier via either one of the two metrics.

In addition, we also exclude cells with exceptionally high *α* values, as inferred by our single-cell HMMs. As cutoff criterion we exclude all cells more than 2 standard deviations higher than the mean value of *α*. This measure tends to overlap strongly with the other outlier criterion (see Fig. S12).

The individual quality control steps are conducted on the level of individual samples and separately per library. The joint copy number analysis is then conducted on the set of cells remaining after application of the replication status and cell-level quality filter. As a consequence of the strict quality control, we end up with a reduced set of high-quality cells. In the case of the TN1-8 ACT samples, we keep between 72% and 80% of input cells after all the above quality control steps.

### Joint segmentation

Any approach estimating single-cell copy number state individually per cell ignores the evolutionary history shared between the cells. Copy number profiles are inherited across cells, apart from the additional CNAs occurring as a consequence of errors during mitosis. Consequently, we can use the shared information to improve the localisation of shared breakpoints and to gain more power to detect copy number changes in individual cells.

In order to combine information across cells, we need to infer the phylogenetic relationship between cells. Here, we describe three different ways to estimate this relationship (step 2 of the algorithm) in the form of a copy number distance matrix *W*. In all cases, the distance is based on patterns of CNAs. The purpose of this work is not in evaluating existing copy number distances, this has been done elsewhere [41], and is beyond the scope of this work. Rather, we demonstrate that *scUnique* can generally support different distance metrics; they all improve the quality of the segmentation. Here, we compare the following ways to estimate *W* : i) the euclidean distance in UMAP space, ii) the distance based on the SCICoNE[35] tree, and iii) the MEDICC2 [41] distance. We refer to the different approaches below as *scUnique*-UMAP, *scUnique*-SCICoNE, and *scUnique*-MEDICC, respectively.

#### UMAP-based distance

In this baseline approach, we identify a neighbourhood of related cells based on a UMAP projection of the cells’ copy number state vectors. We define the distance between two cells *i*_1_ and *i*_2_ based on the euclidean distance in the 2-dimensional UMAP space as 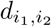. We compute a vector of frequencies of breakpoints across bins for each cell within a percentage of the cell’s closest neighbours (by default 30%). We denote the indicator function of breakpoints obtained in cell *i* at position *j* → *j* + 1 as 𝟙 _i,j_ and [1, *N*_i_] denotes the set of cells belonging to the neighbourhood of cell *i*. We then compute updated per-bin probabilities for a transition of copy number state in cell *i* at position 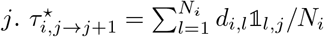

#### SCICoNE-based distance

Here, we replace the UMAP clustering with the phylogeny inferred by SCICoNE. SCICoNE’s phylogeny is based on copy number events, we can directly use the set of cells that share a common history with any given cell as the cell neighbourhood *N*_i_, without using an arbitrary cutoff as in the previous case. We then compute the transition probabilities analogously to the previous case: 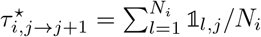

#### MEDICC2-based distance

We use the initial per-cell copy number profiles to infer the phylogeny using the MEDICC2[41] algorithm. MEDICC2 has been specifically developed for use with copy number profiles, and returns a minimal-event distance between copy number profiles.

We use the MEDICC distance matrix *W* to update the time-varying transition matrix entries as follows: 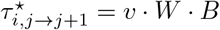 where B is a binary matrix indicating per-cell breakpoints predicted by *scAbsolute-HMM*, and *v* is a weight vector, giving a weight to cells proportional to their read depth *ρ*. The idea being that cells with higher read depths have more reliable breakpoint calls.

It is possible to restrict the cell distances *W* by identifying clusters of cells that are neighbours, equivalent to cutting the phylogenetic tree at a certain level, and so introducing a block structure in the matrix *W*. However, here we opt for the more general solution, in part as determining a good cut-off, or identifying optimal clusters of cells is an open problem.

#### Normalisation

Across all three methods, we normalise the values of *τ*_i,j→j+1_ to a valid probability range, across the entire transition matrix *T*_het_ via a normalisation function ν, with whose help the values of *τ*_i,j→j+1_ are linearly mapped to the range [*τ*_0_, 1 − *ϵ*]:

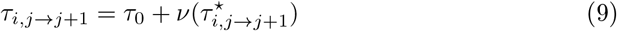

This makes it possible to flexibly adjust the sensitivity of the method in the joint calling step via the value of *τ*_0_. We chose *ϵ* = 1 × 10^−9^, to allow for a very small uncertainty in the transition for the most frequent copy number transition. By default, we use a constant value of *τ*_0_ = 0.05 for *scUnique*, as determined as optimal in the simulation study. *T*_het_ consequently is a diagonal matrix that varies for each transition *j* → *j* + 1.

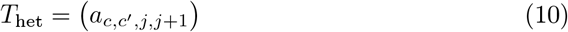

where

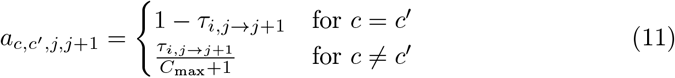

Step 2 of the algorithm relies only on a generic cell distance matrix *W*, and can be extended to other distance measures. We find general improvements in performance when using the information from related cells, and MEDICC2 generally performing best among the variations compared here (see Fig. S13). It is possible to run the inference of *W* (step 2) multiple times, to successively improve the quality of the individual copy number profiles, similar to an EM algorithm. However, this requires re-running the MEDICC2 distance computation which is computationally the most expensive step. The computation of the joint per-cell copy number states is computationally cheap, as we only update the state transition matrix *T* = *T*_het_, and re-run the copy number state inference with the existing parameters. After having obtained a joint copy number segmentation, we then select candidate rCNAs.

### Identifying recent copy number aberrations

We rerun MEDICC2 again on the joint copy number profiles in order to create a final, high quality cell phylogeny. Using this cell phylogeny, we identify the most recent common ancestor in terms of copy number changes for each cell *i*, and define the set of candidate rCNAs as the set of all changes between a cell *i* and its ancestral node in the cell phylogeny. Given the phylogeny, rCNAs are therefore copy number changes that have occurred most recently in the history of the tumour. Note that recentness in this context is not a quantitative property, and we cannot assign a measure of time as to when the changes have occurred.

In order to further reduce the number of false positive rCNAs, we apply multiple post-processing steps to these candidate rCNAs:

Based on the results of the simulation study (Fig. S6), we only select rCNAs with a minimum size (by default larger than 2 bins), or alternatively rCNAs that have a copy number change larger than 2 to include focal, but high amplitude copy number changes. Focal changes are easier to detect because the difference in copy number states is larger, making it easier to distinguish them from random variation observed between neighbouring copy number states.

For all remaining potential rCNAs, we compute the probability of observing the counts given the null hypothesis of the ancestral copy number state. Thus we can directly compare the likelihood of observing the counts under two alternative copy number states and restrict our analysis to a specific event. We compute the following p-value for each segment *S*_l_ running from bin *j*^′^ to *j*^′^ + *k*, assuming knowledge of the correct phylogenetic tree *T*. 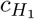 corresponds to the observed change in copy number state under the copy number inference.

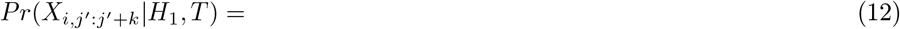

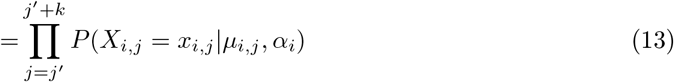

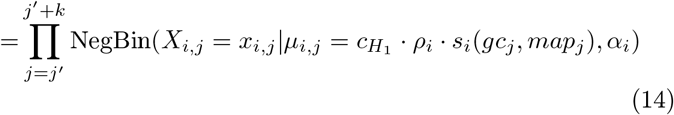

In order to determine the probability distribution under the null hypothesis, we draw *n* = 10 000 samples for every candidate rCNA, assuming the copy number state 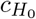 corresponding to the *H*_0_ of the copy number state of the ancestral node, and use this to determine the distribution under the null hypothesis. We correct the resulting p-values using the Benjamini-Hochberg[64] correction, and remove candidate rCNAs that fail a threshold (by default *p*_cutoff_ = 0.01).

Finally, we remove any rCNAs that have identical genomic start and end positions and magnitude. We assume that most of these cases are a consequence of errors in the MEDICC2 phylogeny. It is also possible to apply a more fine-grained filter, e.g. only removing rCNAs with identical start and end coordinates within genomic clones, in the case of heterogeneous samples. In real world data sets, like the TN1-8 patient samples [37], we keep only between 15% and 45% of the putative rCNAs after these strict post-processing steps. This results in a conservative estimate of the number of rCNAs.

### Run time

The run time of *scUnique* is mainly determined by the joint copy number analysis and here in particular the run time of MEDICC2.

For example, running the first part of the workflow – ploidy calling and single-cell HMM inference – takes about 2 h for the TN1 sample (about 1100 cells); it takes about 3.5 h for the 10x Breast A sample (about 2000 cells) and about 14 h for the 10x MKN45 sample (about 5200 cells). These times were measured running the analysis on 64 CPU cores (Intel Xeon E5-2698v4 @2.20GHz), using a 1 Mb bin size resolution. Run time differences between samples are mainly due to the different number of solutions the algorithm has to iterate over in different samples. Importantly, because of the parallel processing of single cells, this part of the analysis has essentially linear run time in the number of cells and can be easily distributed across CPUs.

The joint processing part of the pipeline (run with 32 CPU cores and 156 GB of memory, at 1 Mb bin size) takes similar time to the first step of the workflow for smaller data sets, but scales worse for larger data sets. For example, the run time for the TN1 sample is about 2 h in total (with about 1 h for the initial run of MEDICC2). For the 10x Breast A sample, the total run time is about 5 h, with about 2 h spent on the initial run of MEDICC2. For the 10x MKN45 sample, the total run time is about 40 h (with about 15 h spent on the initial MEDICC2 run). The run time of MEDICC2 itself is determined both by bin size and by the number of cells and is described in detail in Kaufmann et al. [41].

### Simulation study

#### Simulation of copy number profiles

To evaluate the performance of different copy number calling methods on single-cell datasets and to assess the limitations in detecting CNAs on the single-cell level, we conduct a simulation study.

We simulated two distinct types of copy number profiles. The first dataset was designed to represent realistic, relatively uniform copy number profiles. To achieve this, we used 10x Genomics CNV single-cell sequencing data from two ovarian adenocarcinoma cell lines, PEO1 (112 cells) and CIOV1 (339 cells), to generate consensus copy number profiles. Both cell lines exhibit chromosomal instability (CIN) characterised by widespread copy number alterations (CNAs) across the genome.

Per-cell segmentation was performed using QDNAseq [44] with a bin size of 100 kb, corresponding to 27,836 bins. For each bin, we calculated the median copy number across all cells to obtain a consensus profile for each cell line. We chose QDNAseq (a bulk sequencing copy number caller not included in our method comparison) to avoid analytical bias. These consensus profiles served as background single-cell copy number profiles for the simulation of 100 cells for PEO1 and 300 cells for CIOV1.

The second simulated dataset aims to represent a more diverse set of background single-cell copy number profiles, with a more complicated copy number phylogeny that reflects a more heterogenous sample such as a primary tumour. To do this, we use the simulator that is part of the SCICone copy number caller [35] Here we generated a set of 3 background profiles, with 10 000 bins per profile (see Table S2 for the SCICone parameter settings). We took the two sets of background profiles, and then randomly and independently inserted varying numbers (0 to 20 events per cell, from a random uniform distribution) of copy number changes of varying sizes (2, 3, 5, and 10 genomic bins) at random positions on top of these profiles. We simulate changes in copy number state of ± 1 at random, as these are the most challenging CNA to detect. Copy number changes larger than ± 1 are easier to detect due to the greater difference in means between segments.

#### Simulation of reads

Reads are simulated based on the per-cell copy number profiles *y*_i_, using the following formula

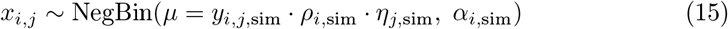

Here, *ρ*_i,sim_ denotes the average reads per bin and per copy number. We vary this parameter, simulating reads for the following values of *ρ*: 3, 5, 10, 15, 20, 30, 40, 50, 100. We use *α* = 0.01 throughout the simulation, as a conservative choice based on what we empirically observe in state-of-the-art scWGS technologies (see Fig. S11). For both *ρ*_i,sim_ and *α*_i,sim_, we draw the per-cell values from a normal distribution with values *ρ*_i,sim_ ~ 𝒩 (*ρ*, 0.25) and *α*_i,sim_ ~ 𝒩 (*α*, 0.002). For cases of *y*_i,sim_ = 0, we draw values from *µ* = 𝒩 (0.3, 0.25) to simulate the presence of non-zero counts that are falsely mapped to certain bins with 0 copy number state. In practice we often observe these small, but non-zero counts in bins that have a zero copy number state, due to some falsely mapped reads and noise.

For all scenarios, we simulate reads with and without the effect of GC content variation. GC variation is simulated via a multiplicative factor *η*_j,sim_ centred around 1.0, which is simulated independently for each cell and varies according to a random uniform variable up to a maximum of approximately 10% from the mean for maxima of GC content. This range covers what we usually observe in practice, when we estimate the impact of GC and mappability content on the mean of the read counts[43]. We set *η*_j,sim_ = 1.0 when we do not simulate GC variation.

#### Method comparison

At the core of identifying rCNAs is accurate copy number calling in scWGS data at the level of individual cells. There is an extensive literature on the copy number segmentation problem. In this study we focus on selected methods that have either been specifically developed or have been adapted and used to analyse scWGS data. Mallory et al. [63] provides a comprehensive review of existing methods and the general workflow for scWGS analysis. Figure 6 gives an overview of existing methods.

**Fig. 6:**
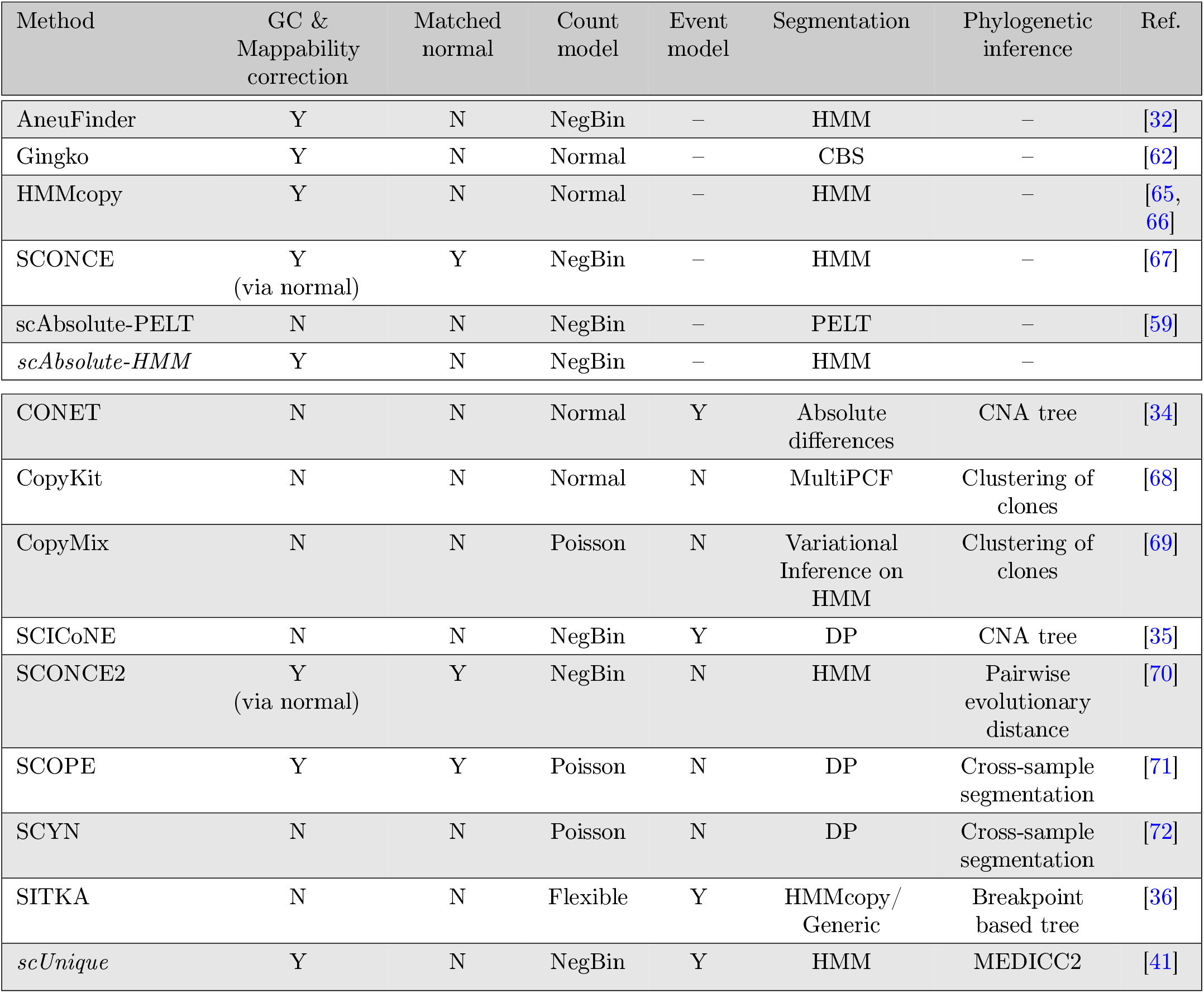
Systematic overview of different copy number calling methods. We distinguish methods focusing on per-cell (top table) and joint segmentation (bottom table). Abbreviations: NegBin: negative binomial, HMM: Hidden Markov Model, CBS: Circular Binary Segmentation, DP: Dynamic Programming.

There are two basic approaches to identify clonal copy number structure. One is to individually segment cells and subsequently determine the cell distances based on their copy number profile [28, 32, 62, 73]. The alternative is to jointly segment and determine cell distances or clusters [34, 35, 37, 71, 74]. The joint approach tends to be computationally more expensive, but offers the advantage of pooling information about breakpoints across cells and so should improve overall quality of copy number calling.

In terms of modelling the noisy count data, methods like QDNAseq [44] and HMM-copy [66] first transform the count data and model it with a Normal approximation, using circular binary segmentation and a HMM for the copy number segmentation, respectively. AneuFinder [32] has specifically been developed for scWGS data under the assumption of a negative binomial distribution.

Here we restrict ourselves to analysing a subset of the methods based on four main criteria: Empirically, we observe that the count data follow a negative binomial distribution, with clear signs of overdispersion (see Fig. S10). Consequently, we restrict our comparison to methods using a negative binomial or Normal likelihood, assuming that any Poisson model would tend to underestimate the variance in the counts, and consequently significantly overestimate the number of rCNAs. In addition, we require the methods to work without using a matched normal sample, or requiring normal cells as part of the input. While it can be very useful to use normal data as a reference for the analysis, and unbiased sequencing can lead to the sequencing of normal cells as a by-product, in many cases normal cells are not available. For one, this excludes many cases of cell lines, where we don’t have a matched normal sample. In addition, even in tumour sequencing experiments, normal cells are not always available, e.g. in cases where the cells have been sorted for certain ploidy levels [37] and sequencing of additional normal cells is relatively expensive. For joint copy number callers, we require the method to generate a form of distance matrix or tree, rather than just clustering the cells and generating clonal level copy number profiles, in order to be able to create a comparison on the level of individual cells. Finally, we require methods to be scalable to thousands of cells in order to be applicable to currently available data sets. The last criterion and the need for a matched sample leads to the exclusion of the SCONCE2 method [70] from our comparison.

Based on these premises, we include the following methods in our analysis: HMM-copy is used as a method that is commonly used in practice [28, 38]. Aneufinder is included as one of the first methods proposing to use a negative binomial count model, albeit segmenting at an individual cell level only. For joint segmentation approaches, we include CONET[34], SCICoNE[35] and SITKA[36], some of the more recently proposed methods. Both CONET and SITKA rely on HMMcopy as input, whereas SCICoNE works independently. It is worth noting that none of the joint methods described above has been developed for the detection of CNAs at the level of individual cells, and consequently, we would not expect any of the methods to perform particularly well at the task.

We compare the performance of individual cell-level segmentation approaches (Aneufinder, HMMcopy, scAbsolute-PELT, *scAbsolute-HMM*) and for joint copy number calling methods (CONET, SCICoNE, SITKA, *scUnique*).

In scAbsolute [43], segmentation and ploidy estimation are performed in two steps. The first step uses a modified PELT algorithm with a negative binomial likelihood to generate an initial segmentation; we refer to this output as scAbsolute-PELT. In the second step, an optional per-cell refinement is carried out by fitting an HMM that leverages the inferred ploidy to produce the final segmentation; we refer to this refined per-cell segmentation as *scAbsolute-HMM*.

Accordingly, in our comparison, scAbsolute-PELT and scAbsolute-HMM represent these two intermediate outputs from the scAbsolute pipeline. Notably, while both stages are part of the same framework, the scAbsolute segmentation differs from the segmentation labelled “PELT” in the original scAbsolute publication.

Note that while the results of the method comparison are dependent on correct ploidy prediction, the results are consistent in cases where ploidy is predicted correctly (e.g. in the SCICoNE simulations). In any case, correct ploidy prediction is a prerequisite for correct copy number estimation in single cells.

We compare performance on two metrics. First, we look at the overall accuracy in copy number calling defined as 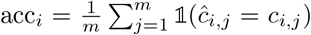, where *c*_i,j_ and ĉ_i,j_ are the simulated and predicted copy number state in cell *i* at position *j*, respectively. In order to keep the results easily comparable, we try to control the impact of ploidy here as far as possible by looking at samples with a ploidy slightly below or around two that should be relatively easy to determine. However, it is important to note, that despite the careful choice of baseline ploidy profiles in the simulations, HMMcopy, Aneufinder, and SCICoNE fail to accurately identify the correct ploidy solution in some cases (particularly in some real world samples). One particular issue we observe is a relationship between the parameters of the segmentation and the resulting ploidy estimates for the samples (see Fig. S7). This makes it difficult to evaluate methods independently of ploidy prediction as described previously [75]. A second issue arising directly from this is the dependency of joint copy number callers such as CONET and SITKA on the results of copy number preprocessing, and HMMcopy, in particular. Initial wrong copy number estimates cascade through the analysis and might contribute to worse overall performance. In the case of SITKA, we address the issue by updating the copy number state between breakpoints with the ground truth ploidy value, e.g. we replace a segment with the “correct”, i.e. majority, copy number state, but keep the predicted breakpoints.

Secondly, we specifically look at the detection of rCNAs. We define genomic regions representing rCNAs as our positive class, and the remainder as the negative class. Thus, we obtain two (highly imbalanced) groups, and we can compute standard metrics such as precision-recall and false discovery rate on this basis. Metrics like the above largely depend on the segmentation threshold, i.e. how sensitive a method is in calling new segments.

In order to allow for a fair method comparison, we initially select the optimal segmentation threshold for each method on two hold-out data sets (PEO1, SCICoNE-2), based on optimal segmentation across a range of values of *ρ* and use the mean F1 score in each data set to select the best parameter across all experiments. We exclude samples whose ploidy is incorrect and thus would negatively bias the selection. We do this for all methods that clearly define a single parameter for their segmentation threshold, such as HMMcopy and SCICoNE. In the case of CONET and SITKA, we use the optimised input from HMMcopy, but there is no single segmentation parameter that could be used to optimise the segmentation, and so we run SITKA with default parameters. In the case of CONET, we run the program using the CBS-mergeLevel segmentation to determine the breakpoints used in the inference process instead of the HMMcopy breakpoint prediction, as recommended by the authors and otherwise default parameters. In the case of *scAbsolute-HMM*, we do not optimise the parameter either, instead we choose a conservative value of *τ*_0_ = 10^−3^. Results for the optimisation of the segmentation threshold for our problem are shown in Fig. S7. For HMMcopy, we vary the parameter ν from 0.1 to 10^4^, for SCICoNE we vary the threshold from − 100 to 40, and for *scUnique* we vary the *τ*_0_ parameter from 10^−4^ to 0.3. The results of the parameter optimisation also indicate that the results are not unduly impacted by the selection of parameters. We chose ν = 100 for HMMcopy, the threshold 30 for SCICoNE, and *τ*_0_ = 0.05 for *scUnique* based on the parameter optimisation. Subsequent analyses are then conducted using the optimal segmentation thresholds for all other data sets (CIOV1, SCICoNE-1, SCICoNE-3). For all other parameters we use the default values. In the case of Aneufinder, we use the method ‘edivisive’, which is the current default option.

#### Pseudobulk copy number signature analysis

Pseudobulk BAM files were generated by aggregating single-cell BAM files using Samtools [76]. Pseudobulk absolute copy number profiles were fitted using 30 kb genomic bins by `swgs-absolutecn` (version 1.3.1) [77]. Copy number signature activity was computed from fitted pseudobulk copy number profiles using `CINSignatureQuantification` (version 1.2.0) [17].

### Experimental methods

#### Cell Lines

The PEO1 cell line, derived from an ovarian adenocarcinoma [78, 79] was cultured in RPMI-1640 supplemented with 10% heat inactivated fetal bovine serum and 2mM sodium pyruvate (Invitrogen) at 37°C, 5% CO_2_. The PEO1 cell line was a gift from Simon Langdon. PEO1 single-cell clones were generated by sorting single cells into 96-well plates containing complete media using the cellenONE® cell dispenser (Scienion). Single-cell clones were cultured for up to 75 days prior to single cell sequencing. The CIOV1 cell line, derived from high-grade serous ovarian carcinoma,[80] was cultured in DMEM/F-12 supplemented with 10% FBS. Cell lines were routinely screened by Phoenix Dx Mycoplasma Mix qPCR (Procomcure Biotech) and tested mycoplasma negative. Cell line identities were authenticated by STR genotyping.

#### Population Doubling Assay

To calculate the population doubling time of the PEO1 clones, for each clone, 1000 cells were seeded into six wells of two 96-Well, Cell Culture-Treated, Flat-Bottom Microplates (Black / Clear Bottom) (Corning ™) in 100*µ*L complete growth media. Six wells were prepared with cell growth media only (Blank). Cells were incubated at 37°C, 5% CO_2_ until assayed. CellTiter-Glo® 2.0 Cell Viability Assay (Promega) was performed at 24 hours post seeding (T(0)), and 7 days post seeding (T(N)), according to manufacturer’s instructions. Luminescence was measured using Clariostar Plus (BMG Labtech). The assay was performed in duplicate using two biological passages.

#### Chromium Single Cell CNV library preparation

PEO1 and CIOV1 cell lines were trypsinised, washed twice with PBS, and counted. The single cell solution was filtered using a 70 *µ*M Flowmi ® cell strainer to remove doublets. 4000 single cells were used for Chromium Single Cell CNV (10x Genomics) library preparation according to the manufacturer’s protocol. Single cell library quantification was performed using the D5000 ScreenTape assay on the Tapestation 4200 platform (Agilent). Samples were multiplexed in equal molarity to achieve 2.4 million reads per cell per sample. Chromium Single Cell CNV constructed libraries were sequenced using the Illumina NovaSeq 6000 S4 platform using PE-150 mode.

#### Cell Cycle Sorting

PEO1, and derived single cell clones, were trypsinised and subjected to methanol fixation and permeabilisation as previously described [43]. For cell-cycle sorting, cells were incubated overnight at 4°C, in Primary Antibody Solution (Geminin E5Q9S XP Rabbit mAb #52508, Cell Signalling Technologies) diluted 1:500 in PBS plus 10% Normal Goat Serum) at a ratio of 100*µ*L per 1 million cells. Cells were then incubated for 1 h at 22°C, in a 1:500 dilution of secondary antibody (Goat anti-Rabbit IgG (H+L) Cross-Adsorbed Secondary Antibody, Alexa Fluor™ 488 #A-11008, Invitrogen). Cells were then incubated in 3*µ*M DAPI solution (1mg/mL, Thermo Scientific) in PBS for 10 min, 1mL per 1 million cells.

To enrich for cells in G1 phase of the cell cycle, cells with negative expression of geminin (expressed only in S and G2/M phase) were gated and sorted using the FACSMelody (BD Biosciences) cell sorter, see Fig. S14 for gating scheme. 435,000 cells were sorted into a 1.5ml Protein Lo-bind Eppendorf tube and subsequently concentrated to 200,000 cells/ml via centrifugation. Single cell isolation was performed using the cellenONE® (Scienion), an image based piezo-acoustic nanolitre dispenser. Details of the cellenONE® selection parameters can be found in Table S4. Single cells were sorted into 384-well plates containing pre-dispensed 300nL 30mM Tris pH8.0 buffer and stored at −20°C until proceeding to library preparation.

#### mDLP+ library preparation

Single cell library preparation was performed using a modified version of DLP+ [28], referred to as mDLP+ [43]. The Mantis® microfluidic liquid dispenser (Formulatrix) and mosquito® HV genomics micropipetter (SPT Labtech) were used for all reagent dispenses and transfers, respectively.

To lyse the single cells, 300nL of freshly prepared 2X protease lysis buffer [prepared from 800*µ*L 2X lysis buffer (30mM Tris pH8.0, 1% Tween-20, 1% Triton X-100) and 200*µ*L QIAGEN Protease (1.36 Anson units/mL prepared in nuclease-free water) was added to each well and incubated for 10 min at 55°C followed by 15 min at 75°C.

For DNA fragmentation and adapter addition, 1.5*µ*L Tagmentation Master Mix (Nextera XT DNA Library Preparation Kit, Illumina) was added per well and incubated at 55°C for 5 min. The master mix was prepared from 1*µ*L TD buffer and 0.5*µ*L ATM buffer per well. The tagmentation reaction was neutralised by 5 min incubation at room temperature with 0.5*µ*L 0.2% SDS buffer.

Generation of dual-indexed single-cell libraries was performed by the addition of 4*µ*L (2X) Equinox Amplification Master Mix (Equinox Library Amplification Kit, Watchmaker Genomics) and 1.5*µ*L 5*µ*M dual-index primers per well, followed by PCR amplification (72°C for 5 min, 98°C for 45s, 12 cycles of (98°C for 15s, 60°C for 30s, 72°C for 45s), followed by 72°C for 5 min, with lid temperature 105°C.)

Primer sequences, as described in [28] were used for dual-index barcoding. 96 individual i7 and i5 primers (Integrated DNA Technologies) were purchased with standard desalting at 100*µ*M concentration in IDTE 8.0 pH buffer. The Bravo NGS Workstation (Agilent) and Mosquito (SPT Labtech) liquid handlers were used to prepare working 384-well plates containing unique dual index primers at 5*µ*M concentration in nuclease-free water containing 0.1% Tween.

Prior to sequencing, libraries were pooled by volume and subjected to two 1X Ampure XP (Beckman Coulter) bead clean-ups. Libraries were analysed using the D5000 ScreenTape Assay for Tapestation 4150 or 4200 (Agilent) and quantified using Qubit 1X dsDNA HS Assay. Libraries were sequenced on the Illumina NovaSeq X using the 25B Reagent Kit (300 cycles) for paired-end sequencing by the Genomics Core Facility at CRUK Cambridge Institute, University of Cambridge. Reads were hard-trimmed and aligned to the human reference genome (hg19) with BWA-MEM using default parameters.

## Data availability

The published single-cell DNA sequencing data used in this study consists of data from three different single-cell whole-genome DNA sequencing technologies.

For 10x Genomics Single Cell CNV datasets, cell lines KATOIII, MKN-45, NCI-N87, SNU-638 and SNU-668 published in [81] are available in the NCBI Sequence Read Archive under accession number PRJNA498809. In addition, three demonstration data sets (a normal diploid Fibroblast cell line (both cell and nuclei), COLO829 G1 sorted 1475 cells, and a Breast Tissue nuclei sections A-E) are published online as part of the 10x Single Cell DNA sequencing technology demonstration (https://www.10xgenomics.com/datasets, see product Single Cell CNV v1.0).

DLP+ data used in this study were published in two separate studies: [28, 38] and are available at https://ega-archive.org/studies/EGAS00001006343, https://ega-archive.org/studies/EGAS00001004448, and https://ega-archive.org/studies/EGAS00001003190. ACT data [37] is available in the NCBI Sequence Read Archive under accession number PRJNA629885 (cell lines BT-20, MB-157, MB-231, MB-453, and tumour samples TN1-TN8).

Single-cell DNA sequencing data from HEK293T ectopic kinetochore experiments [22] are available in the European Nucleotide Archive under accession number PRJEB59859.

In addition, we generated our own experimental data for PEO1 and CIOV1 using 10X Single Cell CNV protocol and PEO1 clones using a modified DLP+ protocol (referred to as mDLP+). These data are available in the European Nucleotide Archive under accession number PRJEB111889.

## Code availability

Using the changepoint package [82] as the basis, we implement our own version of Methods-of-Moments based estimates of negative binomial likelihood for segmentation with the PELT algorithm. The per-cell and joint HMMs are implemented in the Tensor-Flow probability framework [83]. We use MEDICC2 (`2f3934b`) [41] for phylogenetic inference.

The source code for *scUnique* and scripts to reproduce all figures and analyses are available at https://github.com/markowetzlab/scUnique.git. The code used for benchmarking, along with the corresponding software environments and intermediate result files, has been deposited in Zenodo [84]. *scUnique* is using the package environment and genome annotations provided by the QDNAseq package [44] and the ploidy measurements of described in [43].

For the method comparison, we used Aneufinder (v1.20.0), CONET (038b1e4513), HMMcopy (v1.34.0), SCICoNE (50d2e6a629), and SITKA (sitka-nextflow repository, 836b8fe2f7). Software environments for all packages compared are available as part of the simulation code.

## Acknowledgments

We thank all members of Florian Markowetz’s lab for guidance and advice in method development and analysis. In particular we thank Harvey Major and Ang Li. We thank Geoff Macintyre and members of his lab for helpful comments. We thank members of the Brenton Lab, in particular Justina Pangonyte, for assistance with cell line models. We thank Niko Beerenwinkel and the members of the Beerenwinkel lab for helpful discussions and advice on the usage of SCICoNE. We thank Ewa Szczurek and Magda Markowska for helpful advice on the usage of CONET. We thank Sohrab Shah for helpful advice on the DLP+ datasets. We would like to thank the Cancer Research UK Cambridge Institute Bioinformatics, Compliance & Biobanking, Genomics, Flow Cytometry, Histopathology, Microscopy, Research Instrumentation and Cell Services, and Scientific Computing core facilities for their support with various aspects of this study. We specifically thank Ugurcan Sal and Johanna Barbieri for assistance in single-cell sequencing and Jon Marshall for computing support.

## Author contributions

MPS: Method development; MPS, SS: Method implementation and data analysis; DLC: Method development (advice on GAM models); PS: Data analysis (pseudobulk copy number signatures); AEC,: Experimental design and development; AEC, AO, AP: Experimental data collection; MPS, SS, AEC, FM: Manuscript writing; TB, SS: Method and code reproducibility; JDB: Project oversight; FM: Funding, project oversight;

## Supplementary material

**Table S1:** Key varying simulation parameters for SCICoNE. Ploidy, number of cells, and number of bins are identical for the three runs: ploidy = 2, cells = 400, bins = 10000. Explanation of the parameters can be found in the SCICoNE GitHub page.

| Dataset | n regions | n nodes | n reads | Nu | Min Region Size | Max Regions per Node |
| --- | --- | --- | --- | --- | --- | --- |
| SCICoNE-1 | 15 | 100 | 100000 | 10.0 | 2 | 10 |
| SCICoNE-2 | 25 | 150 | 100000 | 4.0 | 5 | 15 |
| SCICoNE-3 | 40 | 20 | 40000 | 4.0 | 1 | - |

**Table S2:**
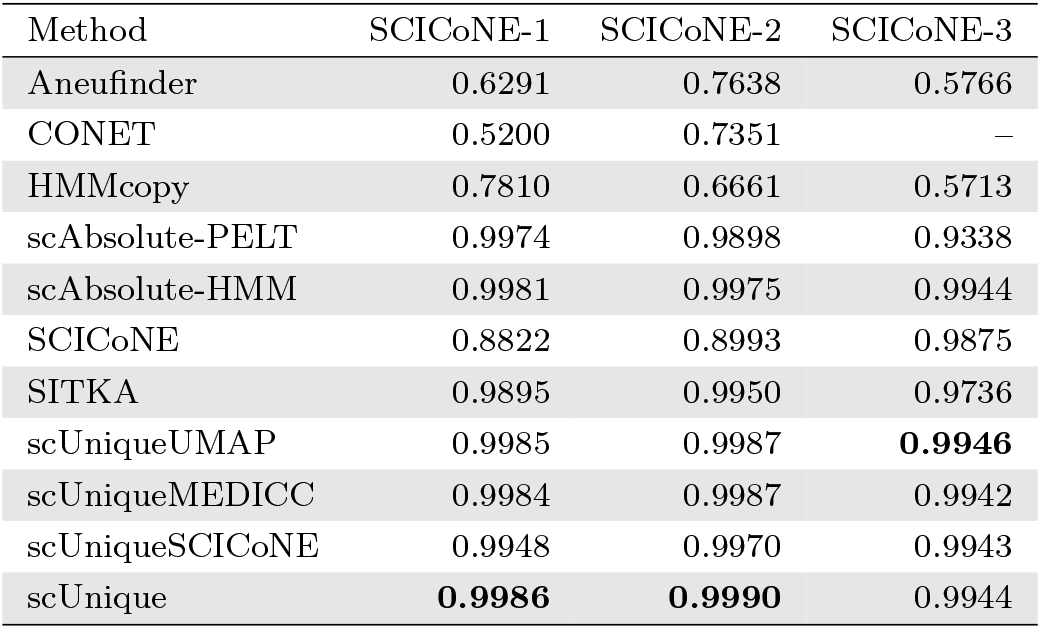
Copy number calling accuracy (defined as the fraction of bins with correctly predicted copy number state, see Methods) on three different single-cell sequencing data sets simulated with the SCICoNE simulator. Note that we did not modify the read counts according to our model, but kept the original data as simulated in SCICoNE. CONET does not converge in the case of SCICoNE-3 within the given limit of iterations. Best performing model is in bold.

**Table S3:** Clone-level doublings, evolutionary position, and filtered rCNA burden.

| Clone | Root Tree Depth | Doublings | Median rCNAs/cell (freq $\leq 5$) |
| --- | --- | --- | --- |
| D7 | 67 | 2.14 | 4 |
| B4 | 58 | 2.63 | 7 |
| C9 | 66 | 2.54 | 8 |
| D5 | 72 | 2.65 | 8 |
| D11 | 129 | 2.66 | 37 |
| G3 | 134 | 2.94 | 48 |

**Table S4:** CellenONE isolation parameters. The cellenONE transmission channel was used to sort single cells based on cell diameter and elongation. These parameters were determined on a individual sample basis using the cellenONE analysis mode. The cellenONE blue channel was used for DAPI gating, and the green channel was used for Geminin-AF488 gating. Abbreviations: CSM, Channel Selection Mode.

| Sample ID | Clone | Min Isolation Diameter ($\mu\text{M}$) | Max Isolation Diameter ($\mu\text{M}$) | Max Elongation | Transmission CSM; Threshold | Blue CSM; Threshold | Green CSM; Threshold |
| --- | --- | --- | --- | --- | --- | --- | --- |
| SINCEL-274 | Parent | 15 | 20 | 1.8 | Positive; 1-255 | Positive; 5-255 | Inspect; 6-255 |
| SINCEL-277 | D11 | 17 | 21 | 1.8 | Positive; 1-255 | Positive; 5-255 | Off |
| SINCEL-279 | B4 | 14 | 18 | 1.8 | Positive; 1-255 | Positive; 5-255 | Off |
| SINCEL-280 | C9 | 14 | 22 | 1.8 | Positive; 1-255 | Positive; 5-255 | Negative; 6-255 |
| SINCEL-281 | G3 | 16 | 22 | 1.8 | Positive; 1-255 | Positive; 5-255 | Negative; 6-255 |
| SINCEL-282 | D5 | 15 | 21 | 1.8 | Positive; 1-255 | Positive; 5-255 | Negative; 6-255 |
| SINCEL-283 | D7 | 18 | 22 | 1.8 | Positive; 1-255 | Positive; 5-255 | Negative; 6-255 |

**Fig. S1:**
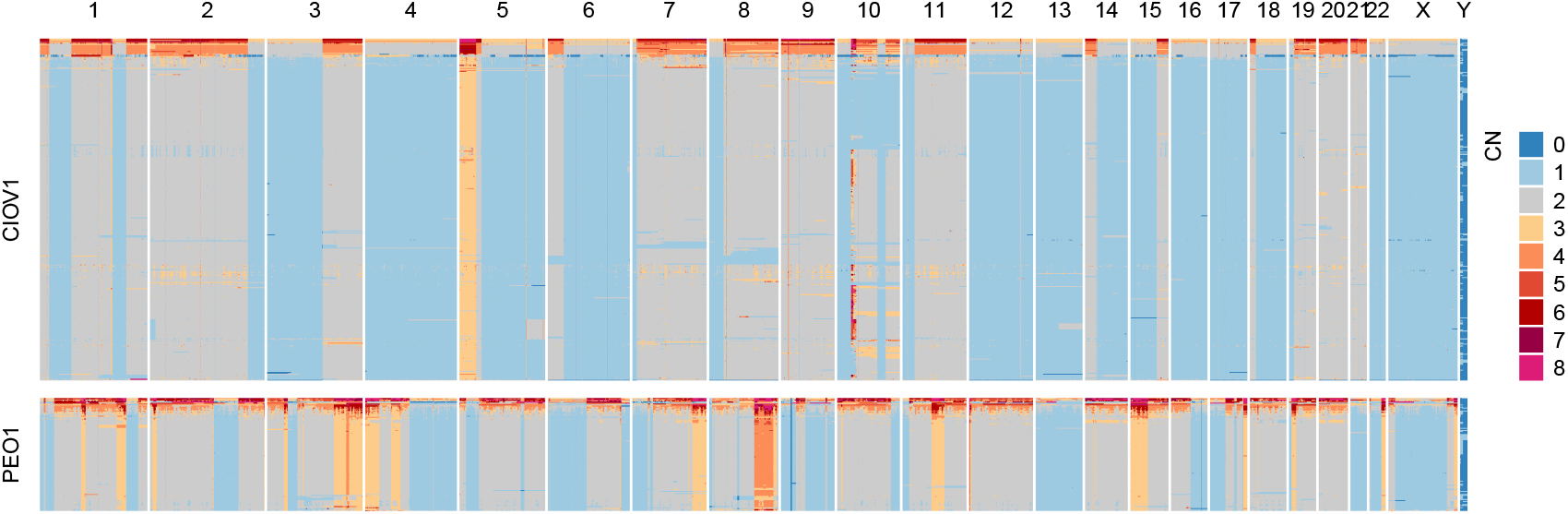
Copy number profiles of PEO1 and CIOV1 cell lines. Heatmaps showing genome-wide copy number profiles for (A) CIOV1 ovarian cancer cells (339 cells) and (B) PEO1 ovarian cancer cells (112 cells). Each row represents a single cell, and columns represent 100 kb genomic bins across all chromosomes (chr 1 22, X, Y). Copy number values are displayed using a colour scale from blue (copy number 0) to red (copy number ≥ 8), with chromosomes separated by vertical lines. Cells are clustered by copy number similarity within each dataset. Data was processed using QDNAseq with 100 kb binning and it was used for making consensus for simulated data.

**Fig. S2:**
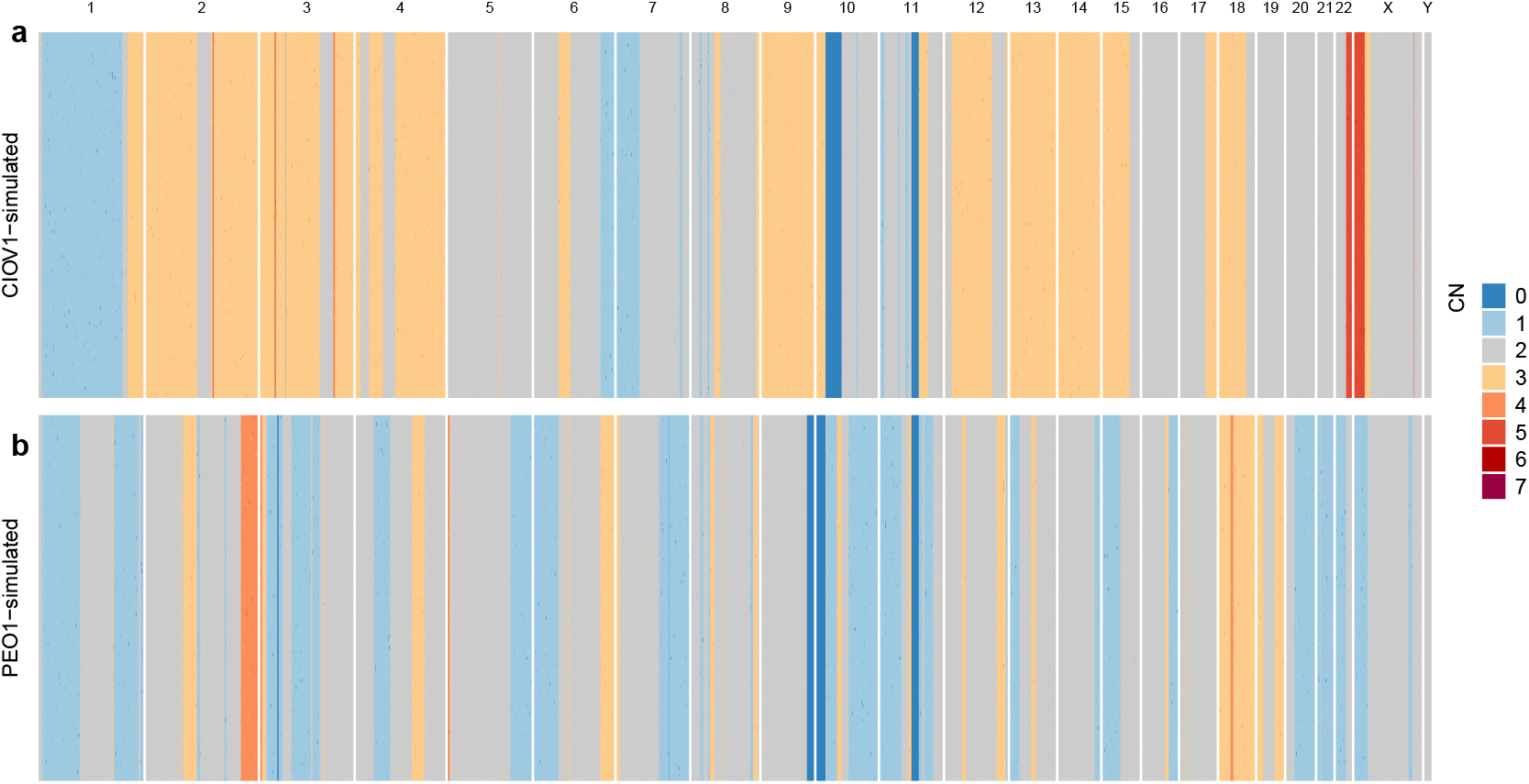
Ground truth copy number profiles for simulated cell line datasets each containing 100 cells. **(a)** CIOV1-simulated dataset. **(b)** PEO1-simulated dataset. For each cell line, we generated a consensus copy number profile from real cells, then simulated individual random CNAs of varying sizes (2, 3, 5, and 10 bins) on top of this consensus to create the per-cell profiles shown. Colours represent integer copy number states ranging from 0 (homozygous deletion) to 7 (high-level amplification), with diploid state (CN=2) shown in gray.

**Fig. S3:**
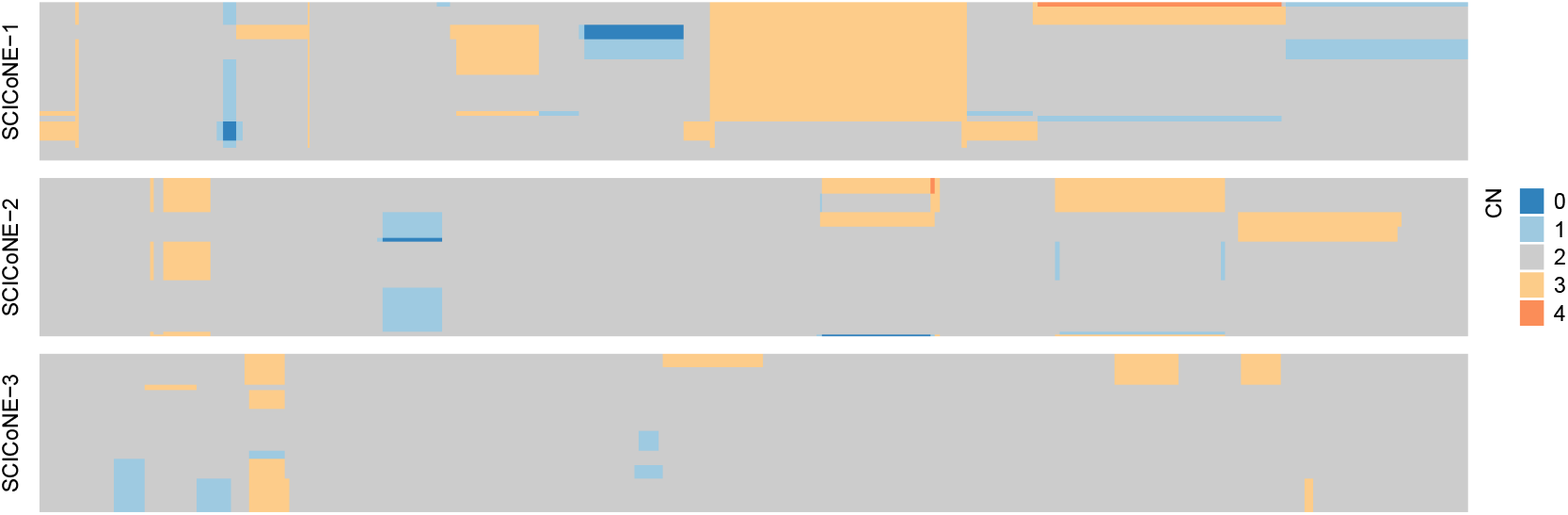
Simulated copy number profiles from SCICoNE. Heatmaps showing simulated copy number profiles generated by SCICoNE for benchmarking purposes. Each panel shows 400 simulated cells across 10,000 genomic bins. Copy number values range from 0-4 (simulations 1-2) or 1-3 (simulation 3), displayed using a colour scale from blue (low copy number) to red (high copy number).

**Fig. S4:**
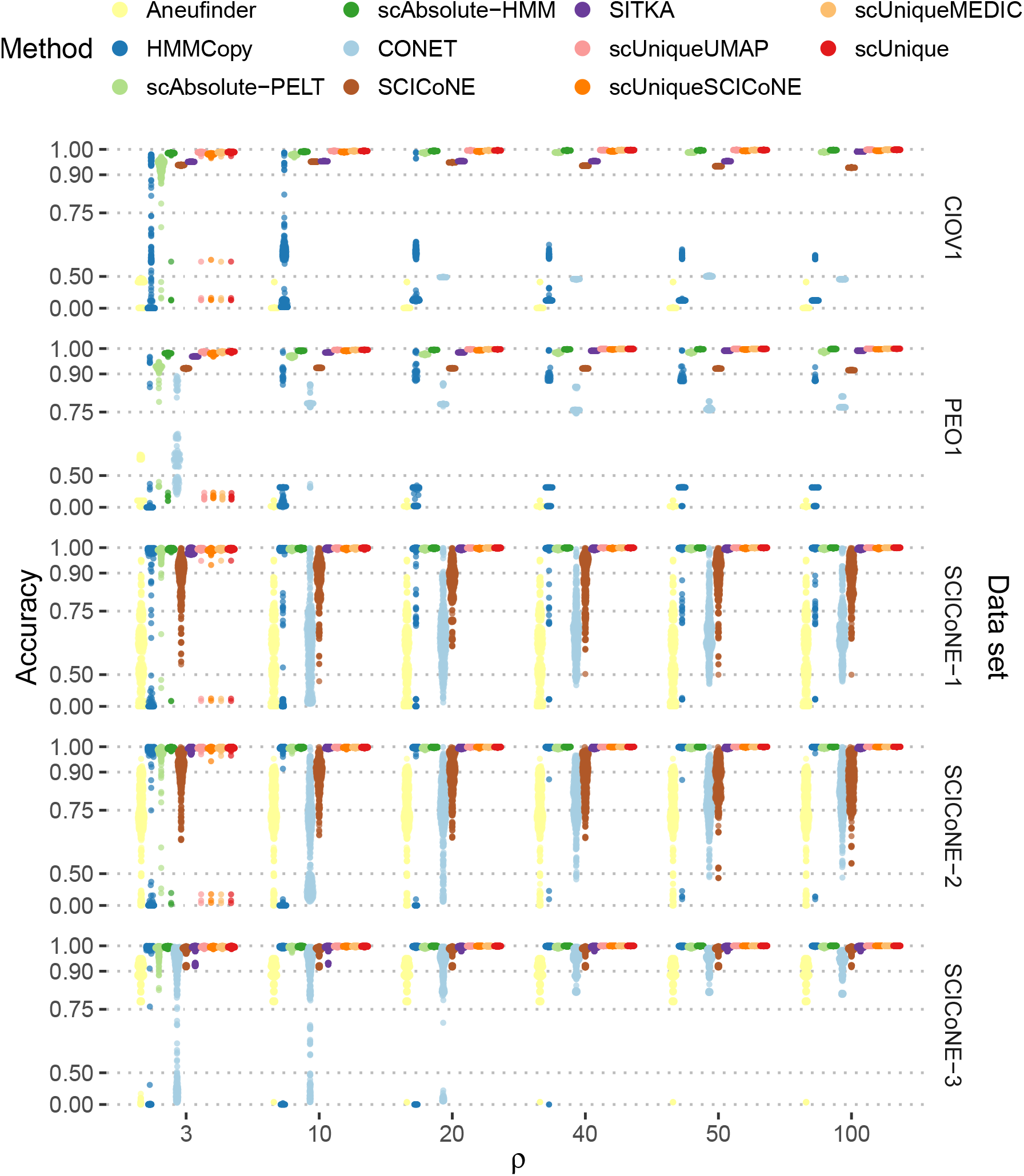
Copy number calling accuracy across five simulated data sets for varying read depth. Overview over overall copy number calling accuracy, for simulations with simulated GC variation. Each data points corresponds to an individual cell. Note that we show only a subset of *ρ* values for better readability. Likewise, the y-axis is on a non-linear scale to better illustrate the differences between better-performing methods.

**Fig. S5:**
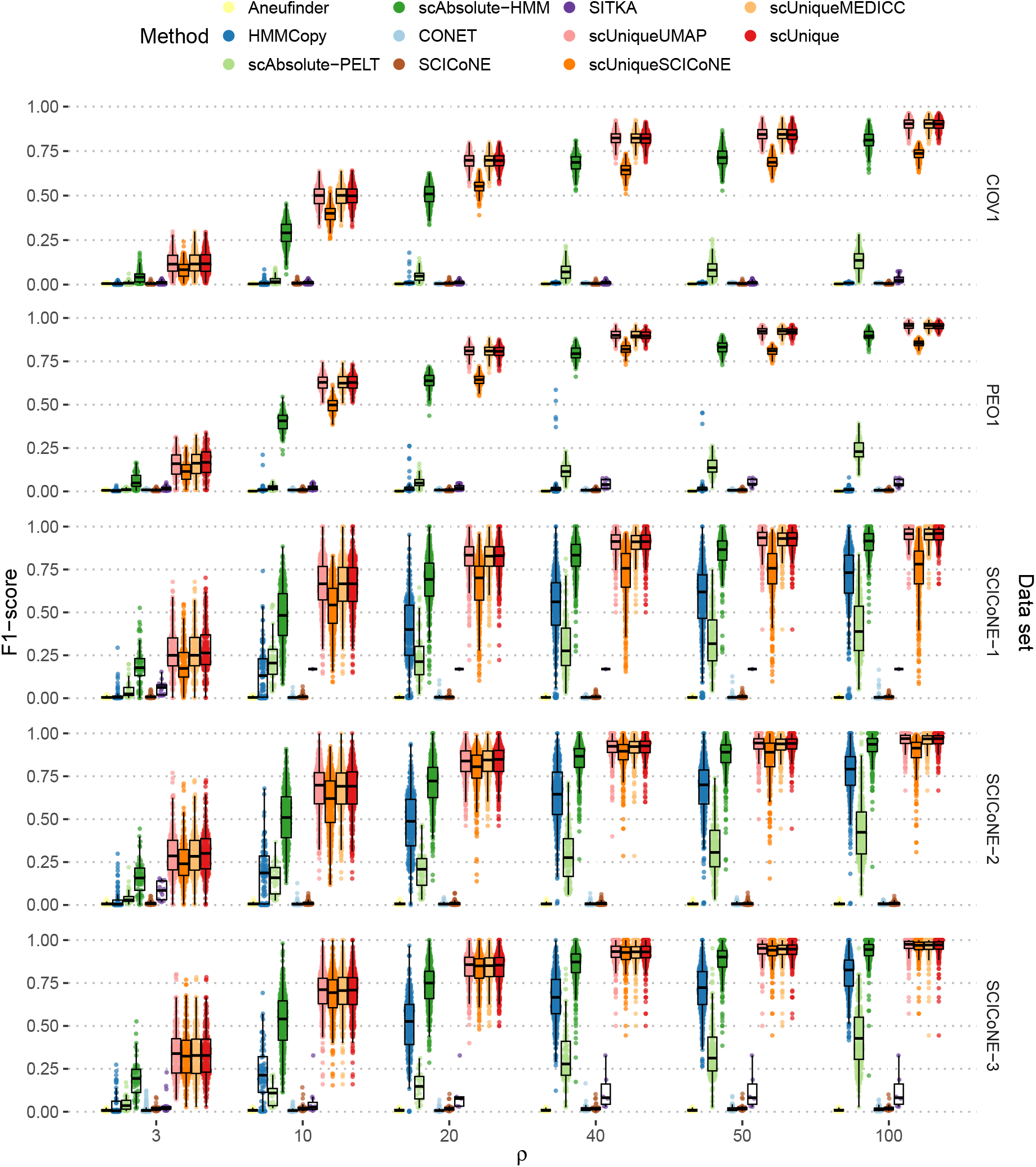
F1 score for detection of rCNAs across five simulated data sets for varying read depth. F1 score, for simulations with simulated GC variation. Each data point corresponds to an individual cell.

**Fig. S6:**
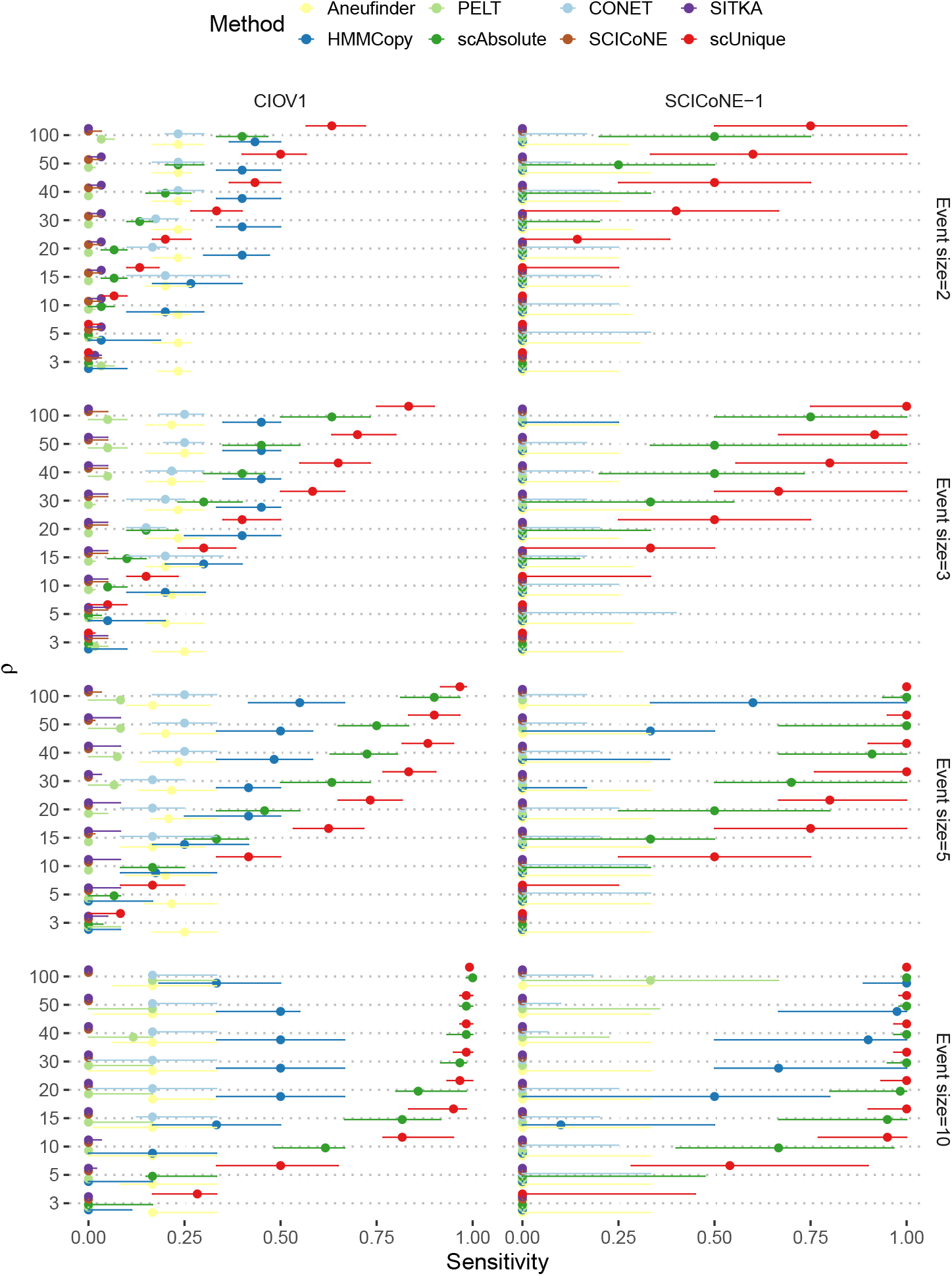
Sensitivity of detecting rCNAs as a function of size of rCNA. Sensitivity for detecting rCNAs in two data sets (CIOV1, SCICoNE-1) for various sizes of rCNAs (2, 3, 5, and 10 genomic bins) is shown. Each point represents the median sensitivity across cells, while the horizontal line extending from the point indicates the 25th and 75th percentiles, respectively. Generally, detection becomes easier with larger event size and higher read depth. For rCNAs of size two, sensitivity is generally low, even for high read depth. Large rCNAs of size ten, however, can be detected by *scUnique* even for lower values of *ρ* around 10. Intermediate-sized rCNAs can be detected at sufficient read depth only by *scUnique* and also *scAbsolute-HMM*, albeit to a lesser extent.

**Fig. S7:**
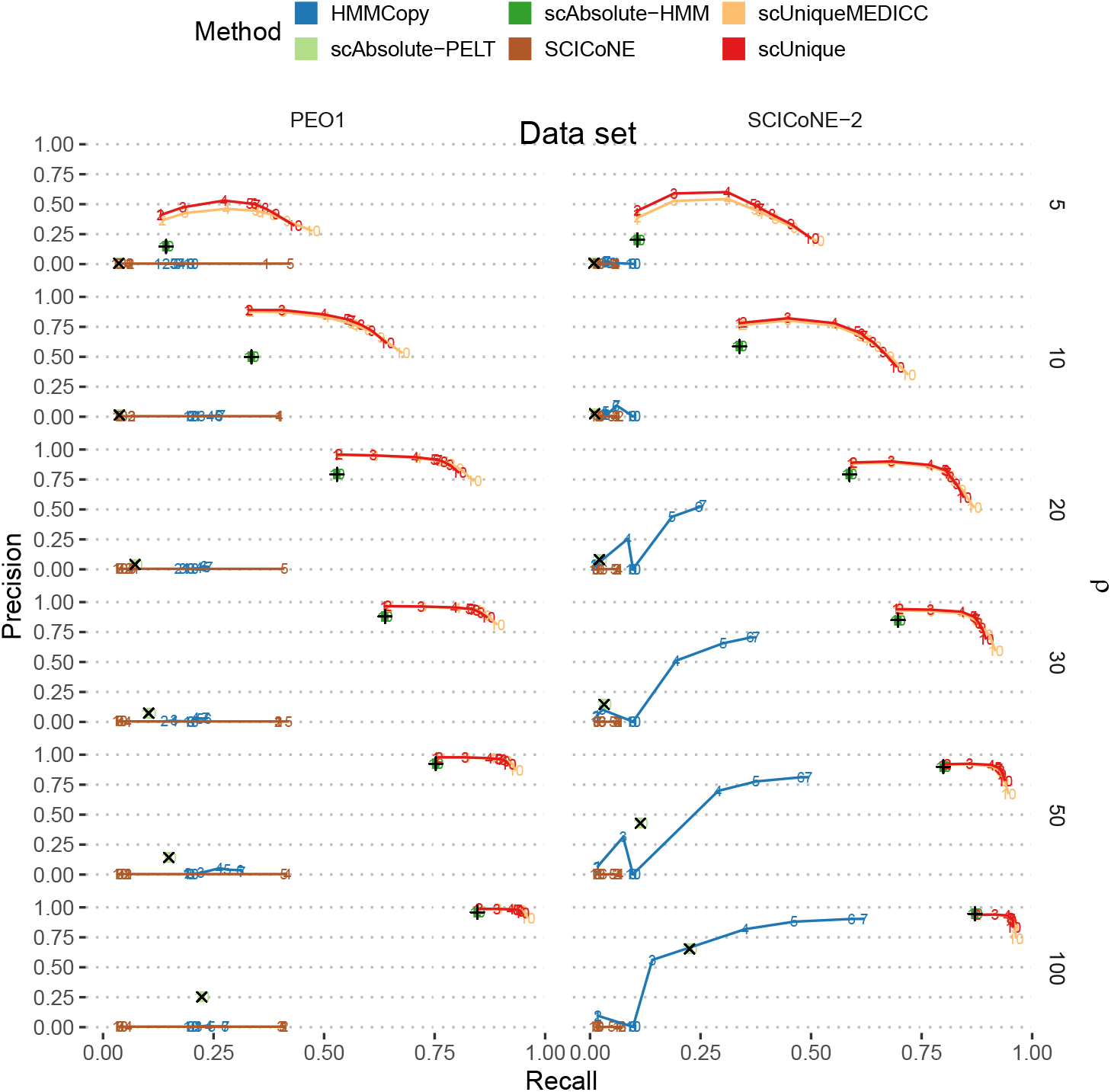
Precision-recall curves for the detection of rCNAs in simulated data sets. A single parameter controlling the segmentation threshold is systematically varied across 10 levels for selected methods in two simulated data sets (PEO1, SCICoNE-2); the number indicates the level of the parameter. Note that the parameter values are different for each method, but have been varied to cover the range of the parameter space. We also include scAbsolute-PELT (light green ×) and *scAbsolute-HMM* (green +) using the default parameter for reference, without tuning their respective segmentation threshold for performance.

**Fig. S8:**
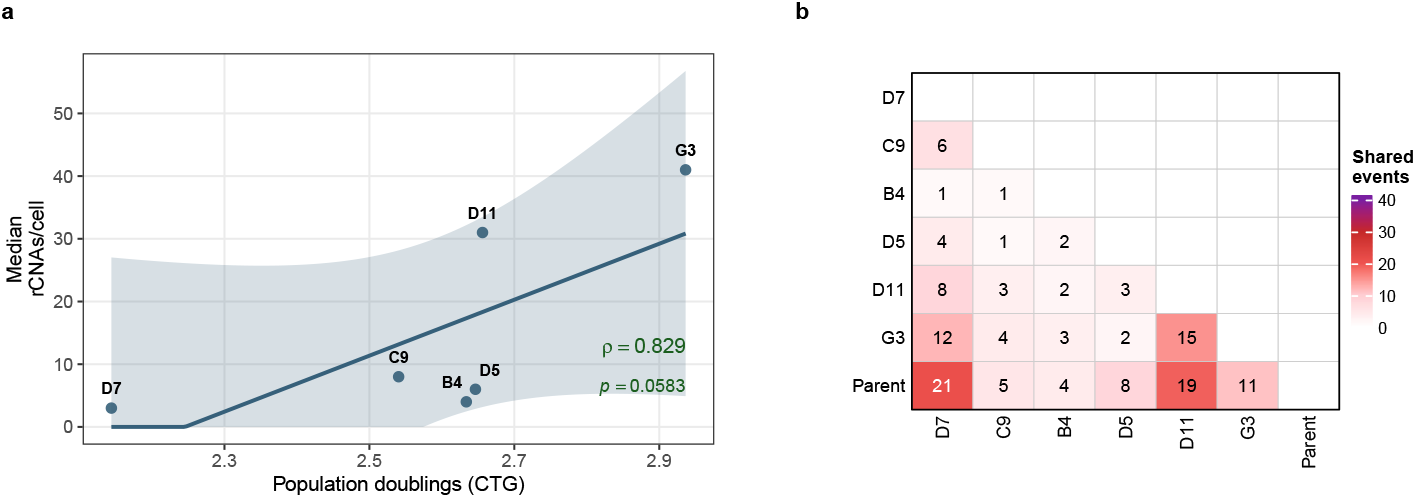
Clone-level rCNA burden and pairwise event sharing across PEO1 lineages. **(a)** Association between population doublings and median rCNAs per cell across clones (Spearman *ρ* = 0.829, *p* = 0.0583). Points represent individual clones; the solid line shows the linear fit and the shaded band indicates the 95% confidence interval. **(b)** Sample to sample sharing matrix showing the number of shared rCNAs between each pair of samples (lower triangle shown). The parent shows highest sharing with D7 (21 events) and D11 (19 events), whereas inter-clone sharing is generally lower (typically 1–15 events), consistent with largely independent post-isolation evolution.

**Fig. S9:**
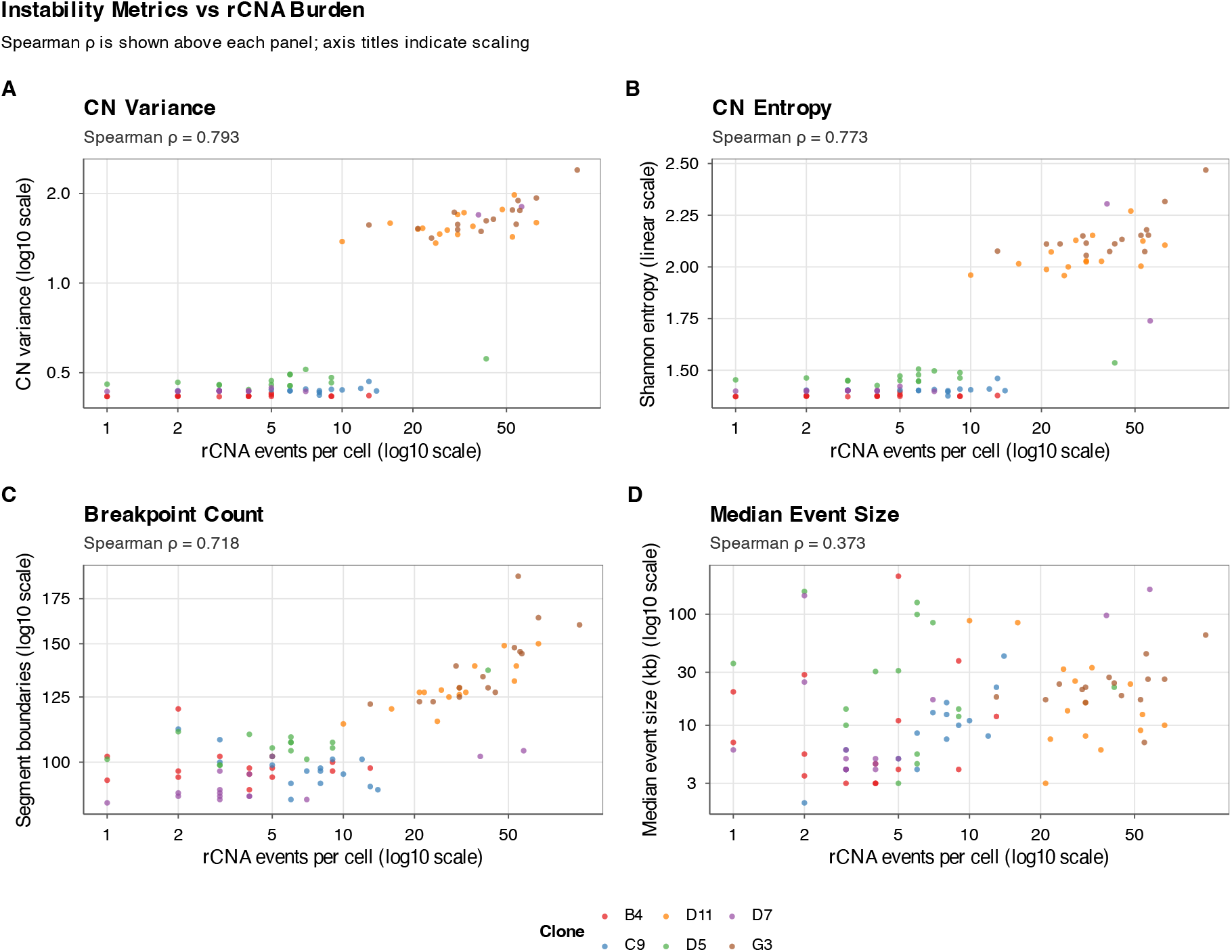
Spearman correlation of rCNA burden with independent chromosomal instability metrics. Scatter plots showing the relationship between per-cell rCNA counts (frequency ≤ 1) and four independent measures of chromosomal instability across six PEO1 clones (B4, C9, D5, D7, D11, G3). Each point represents a single cell, coloured by clone identity. **(A)** Copy number variance shows strong positive correlation with rCNA burden (*r* = 0.775). **(B)** Shannon entropy of copy number states correlates with rCNA counts (*ρ* = 0.759), indicating cells with more rCNAs have more heterogeneous copy number profiles. **(C)** Segment boundary count (break-points) increases with rCNA burden (*ρ* = 0.735), consistent with ongoing genomic rearrangements. **(D)** Median rCNA event size shows weak correlation (*ρ* = 0.324), suggesting event size is largely independent of overall instability burden. These correlations validate that rCNA counts reflect genuine chromosomal instability captured by orthogonal metrics.

**Fig. S10:**
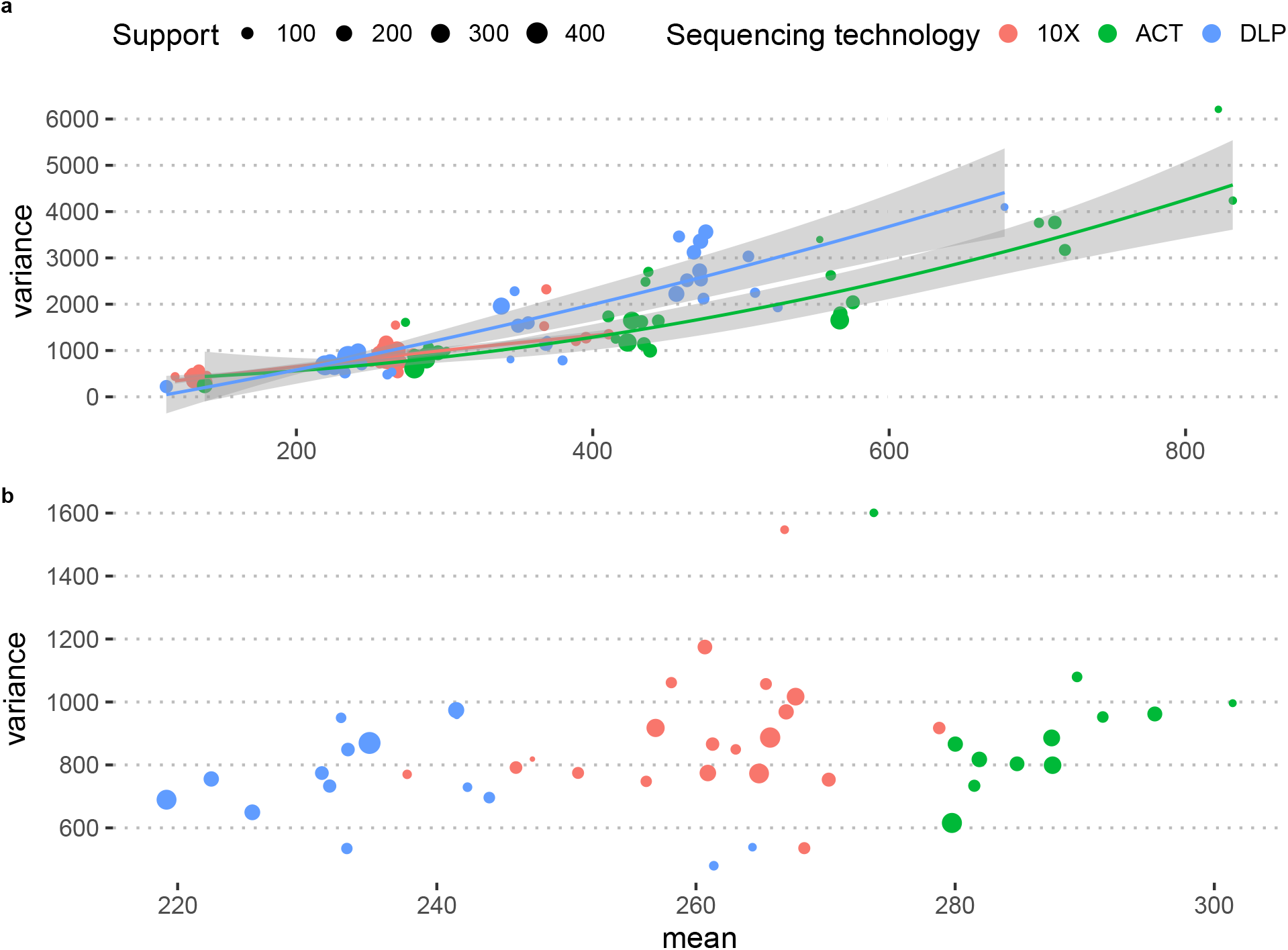
Relationship between mean and variance of raw read counts across the bins in each segment. Data for three example cells, sequenced each with a different scWGS technology (10x, ACT, DLP) are shown. Each point corresponds to a copy number segment, a continuous stretch of bins on the same chromosome sharing the same copy number state. Support indicates the number of genomic bins that each segment is comprised of. We observe that the variance is substantially higher than the mean, indicating that a statistical model taking this overdispersion into account – such as the negative binomial – is required in order to model the mean variance relationship. **a)** Mean-variance relationship across the bins in each segment for all copy number states. **b)** Mean-variance relationship for all counts seen in the bins in each segment with with a copy number state of 2.

**Fig. S11:**
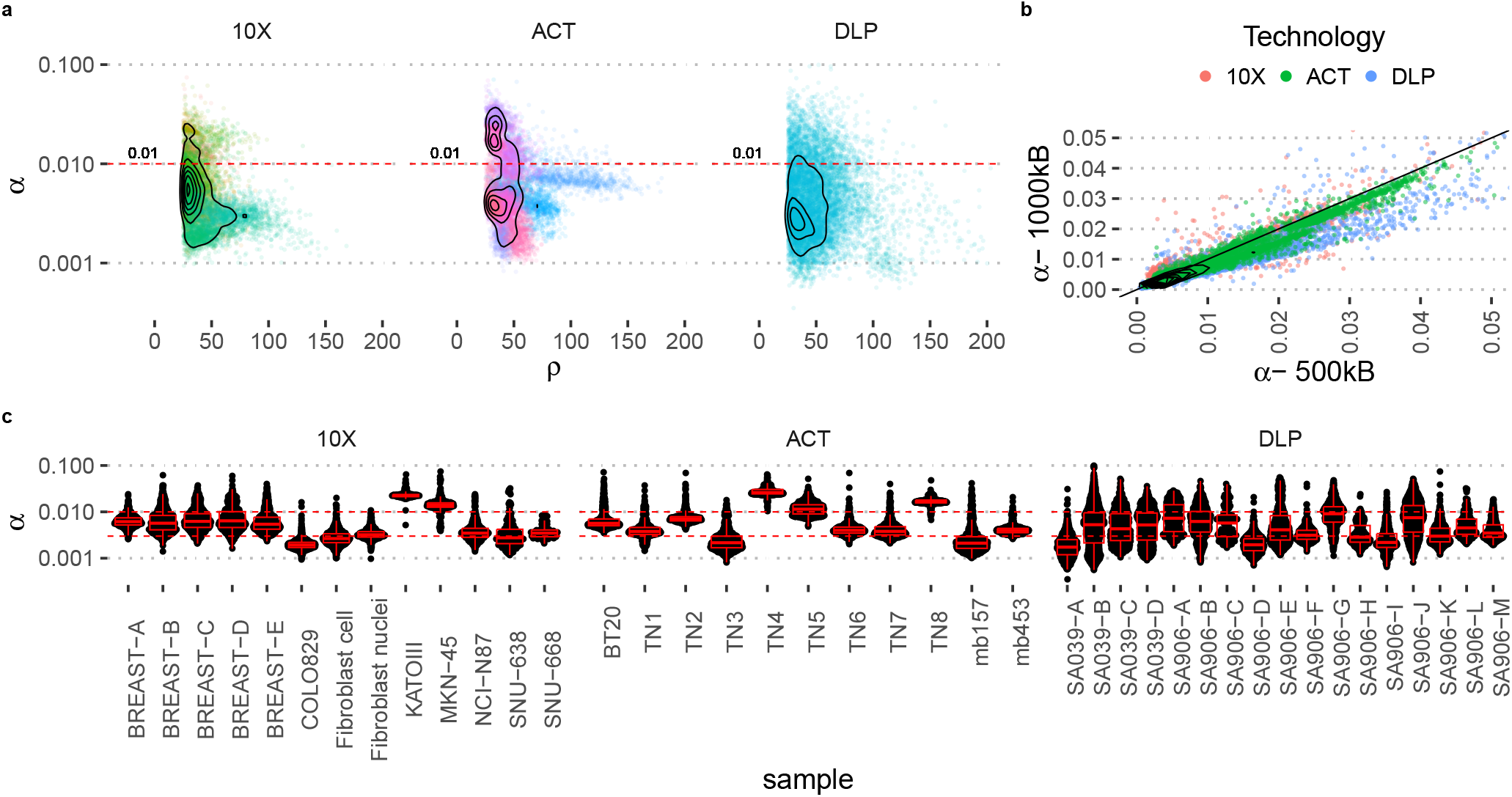
Empirical values of overdispersion *α* across different sequencing technologies. *α* is estimated via the overdispersion parameter of the single-cell HMMs for a bin size of 500 kb. We compare a total of about 10 000 cells for each of the technologies. **a)** Relationship between *α* and read depth *ρ. α* appears to be largely independent of read depth *ρ*. We chose *α* = 0.01 as a conservative threshold for the simulations in this study, indicated by a dashed red line. **b)** Consistency of *α* estimates for bin sizes of 500 kb (x-axis) and 1 Mb (y-axis). **c)** Technical variation in *α* across samples and sequencing libraries. Generally most samples fall within a window of *α* = 0.01 to *α* = 0.003, indicated by the two dashed, horizontal red lines.

**Fig. S12:**
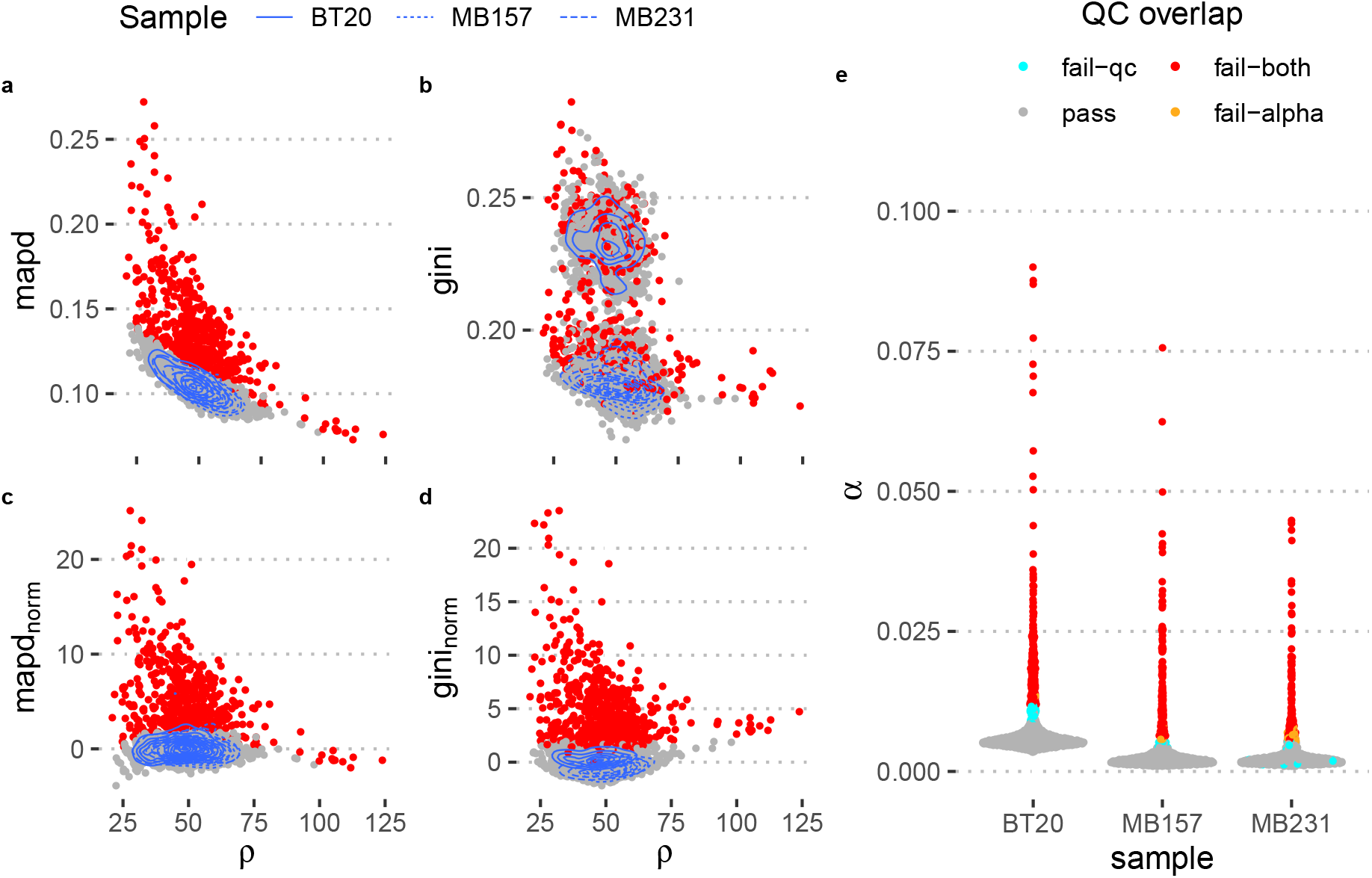
Per-cell quality metrics before and after normalisation. Quality metrics for three different cell lines (BT20, MB157, MB231) sequenced with ACT technology. We use the cell lines as a proxy for a heterogeneous cell population. Colours indicate cells that have been excluded as consequence of (red) or passed (grey) the QC process. Density plots (blue) indicate areas of highest density for the three data sets corresponding to three different cell lines (indicated by line type). **a)** and **b)** show the unnormalised Gini and MAPD coefficients for each cell. **c)** and **d)** show the normalised Gini and MAPD values, where the effect of read depth and copy number profile, respectively, have been removed. **e)** Distribution of *α* values across all three samples and outlier detection based on *α* value. Cells with high values of *α* typically also fail the quality check based on MAPD and Gini coefficient, but there are also cells that are excluded only by the *α* criterion or the quality check. Colours indicate cells that fail either based on the QC criterion (Gini and MAPD, in light blue) or based on the *α* value criterion (orange) or that fail both criteria (red) or that pass QC (grey).

**Fig. S13:**
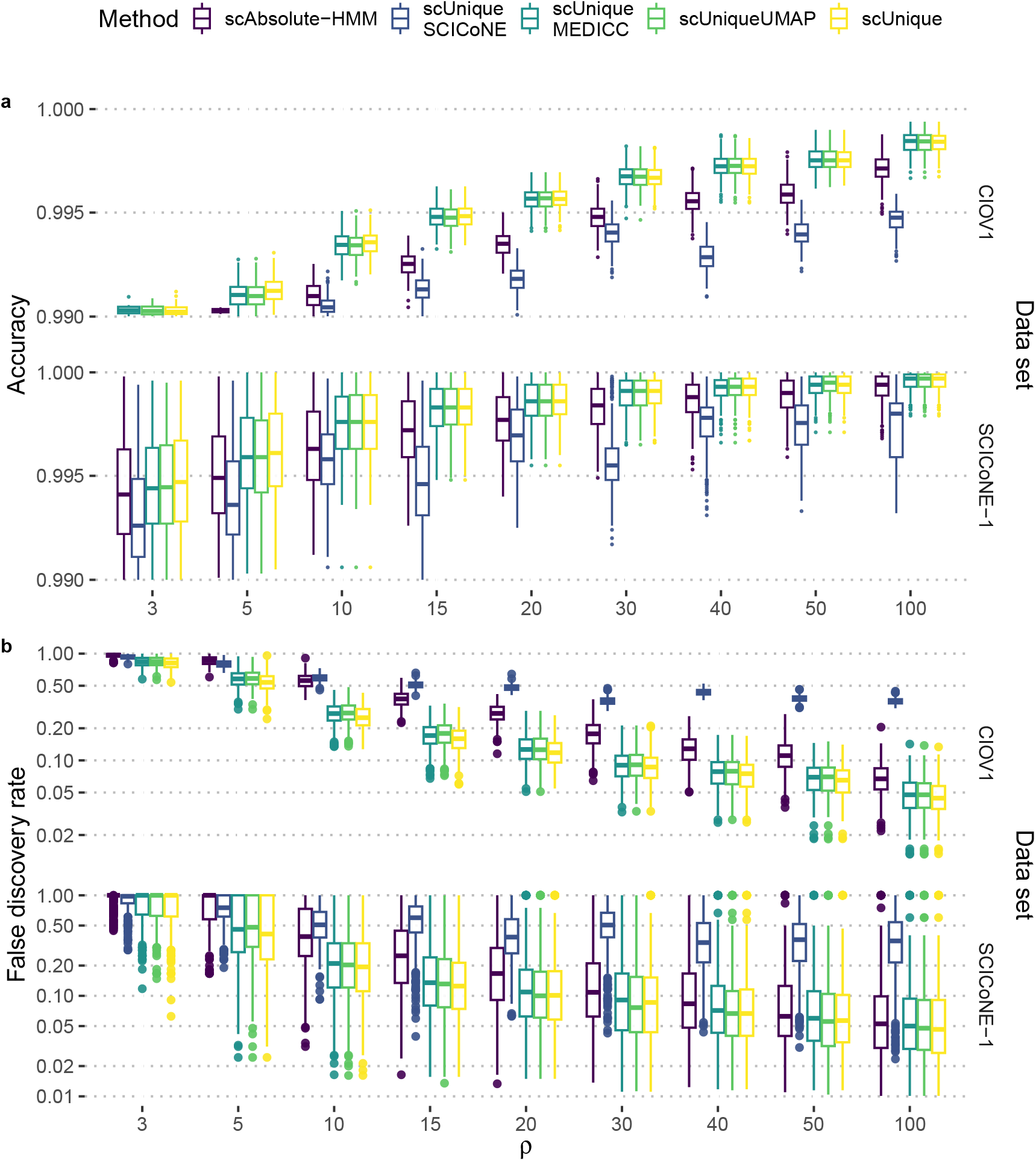
Comparison of accuracy and false discovery rate between different version of the *scUnique* algorithm. Performance is shown for *scAbsolute-HMM*, three different versions of the *scUnique* algorithm that use different ways of estimating the cell-by-cell distance matrix *W* (UMAP, SCICoNE, or MEDICC2), and the complete version of *scUnique* that includes step 3 of the algorithm, i.e. a selection of high-confidence rCNAs only. **a**) Accuracy of copy number calling for two data sets for increasing read depth *ρ*. **b**) False discovery rate for detecting simulated rCNAs for two data sets for increasing read depth *ρ*.

**Fig. S14:**
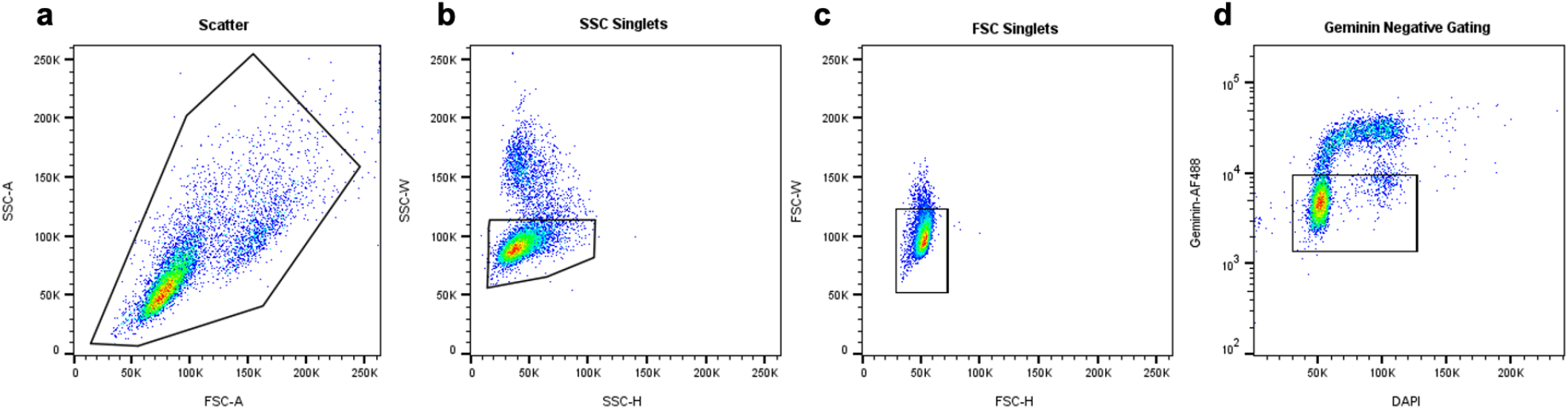
Example gating scheme for G1-phase cell cycle sorting using geminin. Cell lines were methanol fixed and permeabilised prior to incubation with Geminin-AF488 antibodies and DAPI staining. Acquisition, analysis and sorting was performed on the BD FACSMelody using the BD FACSChorus™ software. Analysis was performed using Flow Jo v10.10. PEO1 Clone B4 used to demonstrate gating. The gating tree was as follows: **a**) FSC-A/SSC-A (gating of cells based on size and intracellular composition) **b**) SSC-H/SSC-W and **c**) FSC-H/FSC-W for removal of non-singlets **d**) DAPI-A/Geminin-AF488-A for cell cycle analysis. Geminin is a DNA replication regulator expressed in S and G2 phase of the cell cycle, and not expressed in G1-phase of the cell cycle. DAPI is used as a DNA content marker, and is routinely used in cell-cycle analysis. Together, geminin and DAPI intensity are used here to gate cells in G1-phase of the cell cycle. The use of a primary-antibody negative control population (AF488 and DAPI staining only) was used to determine the gating of cells with negative expression of geminin. The geminin-negative population were sorted via FACS and subsequent single cell isolation was performed using the cellenONE® cell dispenser.

**Fig. S15:**
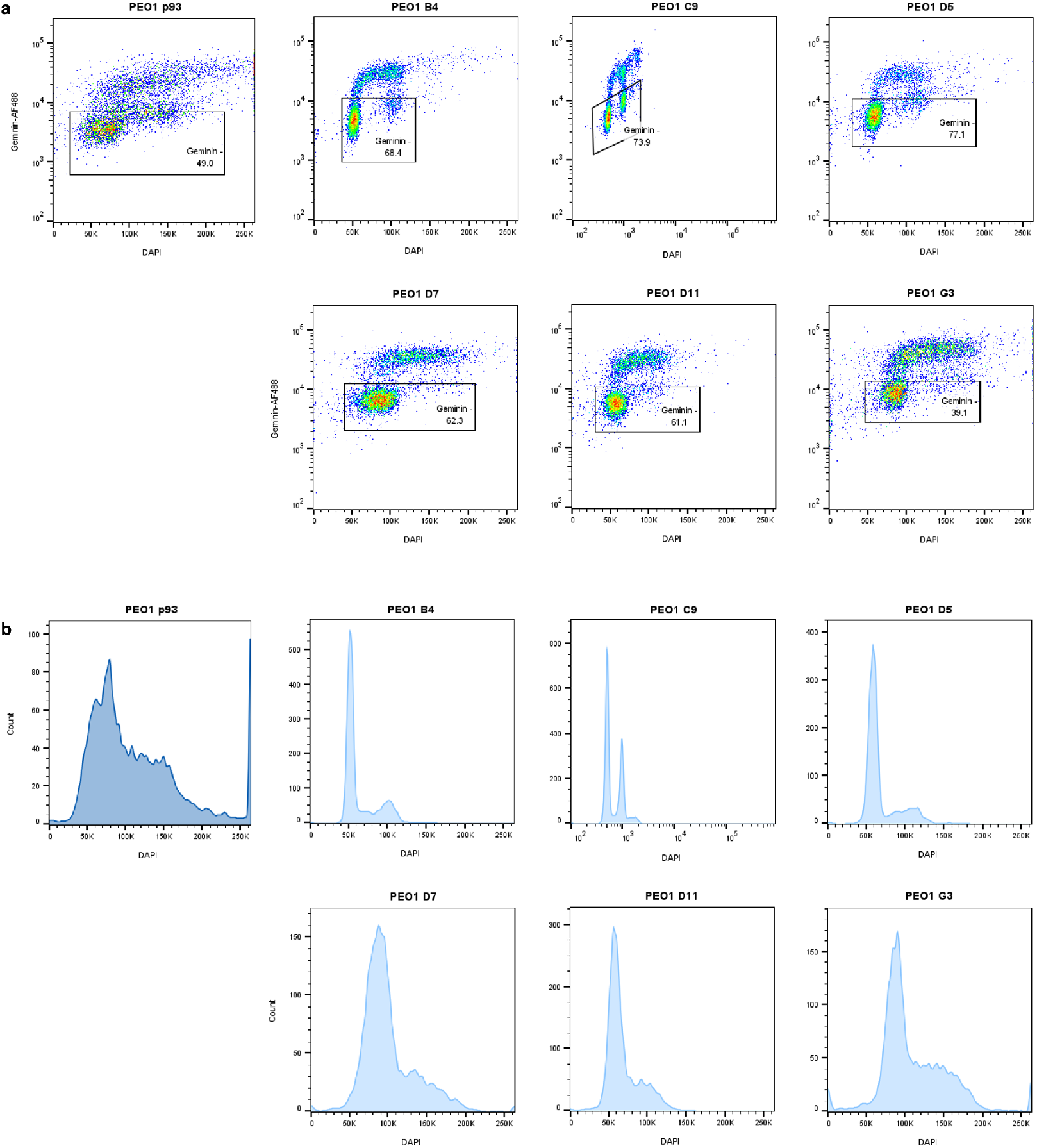
Geminin and DAPI intensities of PEO1 single-cell clones. FACS analysis of PEO1 single-cell clones showing differences in DNA content between the individual clones. Acquisition, analysis and sorting was performed on the BD FACSMelody using BD FACSChorus™ software. Analysis was performed using Flow Jo v10.10. Due to the proportional binding of DAPI to DNA, DAPI intensity is displayed using a linear axis. Note, due to error during data acquisition, PEO1 C9 uses logarithmic scaling on the x-axis. **a**) DAPI/Geminin-AF488 for cell cycle analysis. **b**) DAPI histogram. PEO1 p93, the parent sample, of which the single-cell clones was derived has a broad range of DAPI intensities, indicating a highly heterogenous population with significant aneuploidy, whereas the single-cell clones are more homozygous. PEO1-C9 sees a population of cells that are have a high DAPI intensity, and low Geminin intensity, that suggests a sub-clonal population following a whole doubling event. This is not supported in the single-cell copy number analysis for PEO1-C9 clone. This population could be 1) G2 cells with weak Geminin staining, 2) doublet cells that were not screened out via up-stream FACS gating. 3) an artifact of the logarithmic acquisition of the DAPI channel. The broad peaks seen in D7 and G3 suggests chromosomal instability with individual cells harbouring copy number gains and losses.

